# A neurogenetic mechanism of experience-dependent suppression of aggression

**DOI:** 10.1101/2020.06.26.172890

**Authors:** Kenichi Ishii, Matteo Cortese, Xubo Leng, Maxim N. Shokhirev, Kenta Asahina

## Abstract

Aggression is an ethologically important social behavior^1^ but excessive aggression can be detrimental to animal fitness^2,3^. Social experiences among conspecific individuals reduce aggression in a wide range of animals^4^. However, the genetic and neural basis for the experience-dependent suppression of aggression remains largely unknown. Here we found that *nervy* (*nvy*), a *Drosophila* homolog of vertebrate myeloid translocation gene (MTG)^5^ involved in transcriptional regulation^6–8^, suppresses aggression via its action in a specific subset of neurons. Loss-of-function mutation of the *nvy* gene resulted in hyper-aggressiveness only in socially experienced flies, whereas overexpression of *nvy* suppressed spontaneous aggression in socially naïve flies. The loss-of-function *nvy* mutant exhibited persistent aggression under various contexts in which wild-type flies transition to escape or courtship behaviors. Knockdown of *nvy* in octopaminergic/tyraminergic (OA/TA) neurons increased aggression, phenocopying the *nvy* mutation. We found that a subpopulation of OA/TA cells specifically labeled by *nvy* is required for the social-experience-dependent suppression of aggression. Moreover, cell-type-specific transcriptomics on *nvy*-expressing OA/TA neurons revealed aggression-controlling genes that are likely downstream of *nvy*. Our results are the first to describe the presence of a specific neuronal subpopulation in the central brain that actively suppresses aggression in a social-experience-dependent manner, illuminating the underlying genetic mechanism.

---

Animals adjust their aggressiveness toward conspecifics according to the perceived costs or benefits of the interaction^3^. An animal’s prior social experiences play an important role in either promoting or suppressing the intensity of aggression^4^. Recent studies have focused primarily on neuromodulatory factors and circuits that promote aggression^9–15^. To date, only a handful of genetic and neural substrates have been associated with suppression of animal aggression^16–19^. Here we specifically sought to unravel the neurogenetic mechanisms that mediate experience-dependent suppression of aggressive behavior, a phenomenon widely observed from invertebrates to vertebrates^4^. To this end, we used the fruit fly *Drosophila melanogaster* to identify genes necessary for suppressing aggression under group-rearing conditions, where flies are provided with social experiences that normally reduce aggression. We performed a systematic behavioral screen using adult male flies with candidate genes knocked down in neurons via RNA interference (RNAi). Fly aggressiveness was quantified by automated counting of lunges, a male-type aggressive behavior^20^, performed among male pairs. Among 1,408 RNAi effector lines each driven by the pan-neuronal *elav-GAL4* driver, flies in 11 lines showed a significant increase in lunges compared with genetic controls after group rearing (Fig. 1a). The most prominent increase was produced by neuronal knockdown of the gene *nervy* (*nvy*) (Fig. 1a-b and Extended Data Fig. 1a), which we focus on herein. The heads of the *nvy* RNAi flies showed reduced levels of *nvy* mRNA (less than 40% of controls) and Nvy protein expression (Extended Data Fig. 1b).

**Fig. 1.**
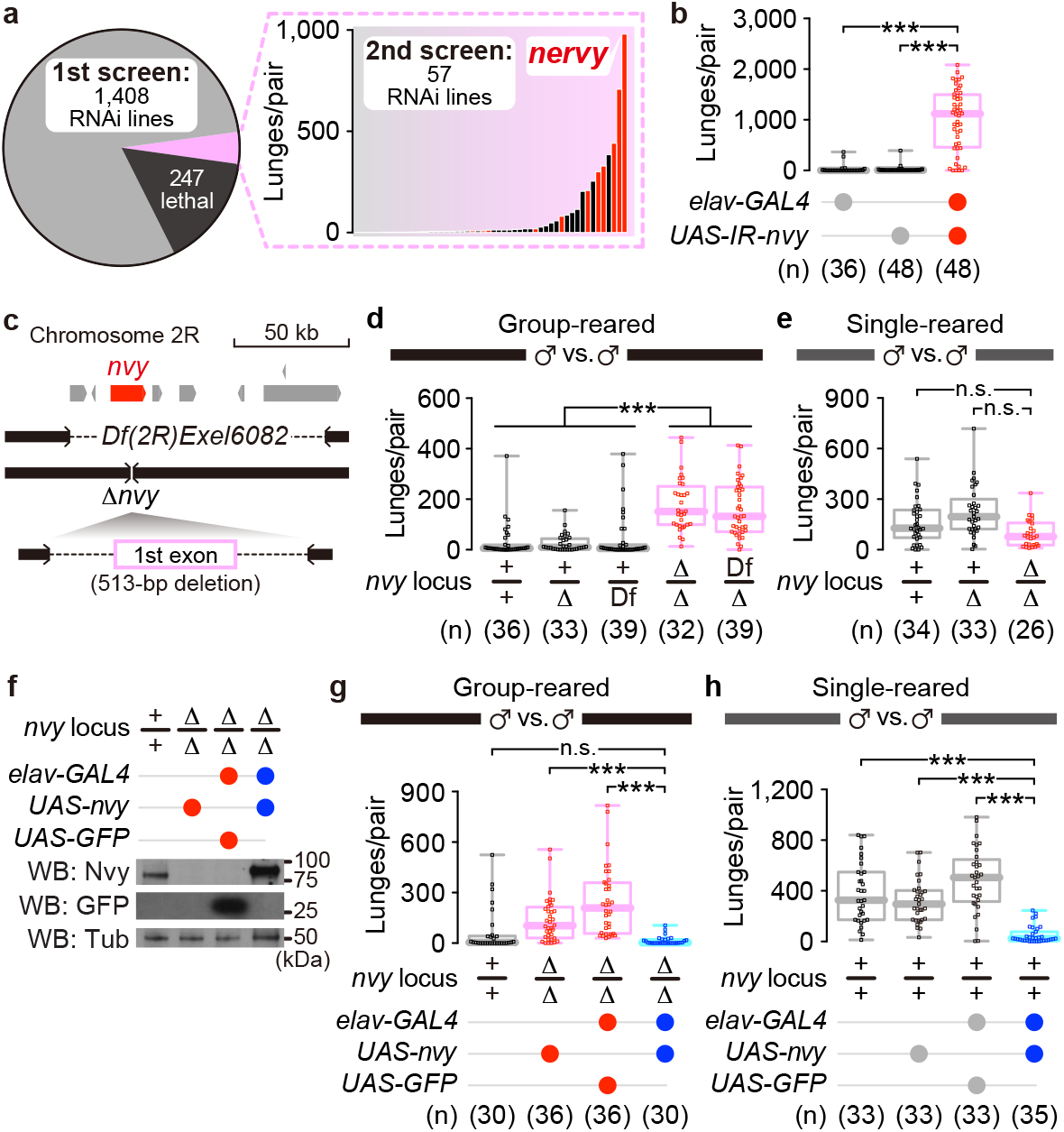
Identification of the *nvy* gene as a negative regulator of aggression. **a**, Results of the pan-neuronal RNAi screen for *Drosophila* genes that suppress aggression. The median number of lunges in 30 min for pairs of knockdown mutants (20–96 pairs for each line) are shown in the bar plot (right). Red bars represent 11 RNAi lines that showed a significant increase in lunges compared with two genetic controls harboring either *elav-GAL4* or *UAS-IR* (Inverted Repeat). For full descriptions of the RNAi screen results, see Supplementary Tables S1-2. **b**, Increase in lunges performed by males with pan-neuronal RNAi of the *nvy* gene, shown in box plots. **c**, Genome schematics of two deletion alleles, a deficiency *Df(2R)Exel6082* and the CRISPR/Cas9-mediated knockout *Δnvy*. Pointed boxes colored in gray and red (*nvy*) represent 8 protein-coding genes disrupted by *Df(2R)Exel6082*. **d**, Increased aggression by deletion of *nvy* in socially experienced flies. For the *nvy* locus genotypes, “Df” and “Δ” represent *Df(2R)Exel6082* and Δ*nvy*, respectively. **e**, Induction of the hyper-aggressive phenotype in Δ*nvy* mutants requires prior social experience. **f**, Pan-neuronal expression of Nvy in the Δ*nvy* background verified by Western blot. α-Tubulin (Tub) was detected as an internal control. **g**, Rescue of the hyper-aggressive phenotype in Δ*nvy* males by the pan-neuronal *nvy* expression. **h**, Reduced aggression by the pan-neuronal *nvy* overexpression. For each box plot, the thick line represents the median, the box extends from the 25^th^ to 75^th^ percentiles, and the whiskers show the range from minimum to maximum. *** p < 0.0005, n.s. p ≥ 0.05; Kruskal-Wallis one-way ANOVA (**b,d,e,g,h**) and post-hoc Mann-Whitney U-test with Bonferroni correction (**b,e,g,h**) or Dunn’s multiple comparisons test (**d**). Flies with the same genotype were paired, and the numbers of tested pairs are shown in parentheses.

We generated a CRISPR/Cas9-mediated null mutation of the *nvy* gene (Δ*nvy*; Fig. 1c) and confirmed that *nvy* is indeed necessary to dampen aggressiveness after group rearing. The homozygous Δ*nvy* mutation, as well as trans-heterozygosity of Δ*nvy* and *Df(2R)Exel6082*, a chromosomal deficiency lacking the 110-kb genomic region that encompasses *nvy*, led to increased aggressiveness relative to genetic controls after being reared in groups of 15 individuals for 5–7 days (Fig. 1d). However, when Δ*nvy* homozygous mutants were reared in isolation, a condition known to elevate basal aggression^21^, they showed similar levels of aggression as wild-type flies (Fig. 1e). These results suggest that *nvy* is involved specifically in social-experience-dependent suppression of aggression. The locomotor activity of flies introduced into the arena solitarily was comparable across genotypes (Extended Data Fig. 1c), arguing against the possibility that the high level of aggression in Δ*nvy* is due to general hyperactivity. Pan-neuronal *nvy* expression *via* the *UAS-nvy* transgene (Fig. 1f) reversed the hyper-aggressive phenotype of the group-reared Δ*nvy* mutants (Fig. 1g). In addition, *nvy* overexpression in the wild-type background reduced basal aggressiveness in single-reared flies (Fig. 1h) without reducing their locomotion (Extended Data Fig. 1e). These results establish *nvy* as the first gene known to be both necessary and sufficient to suppress aggression under group-rearing conditions.

*nvy* (named after its abundant expression in the nervous system^5^) was identified as a *Drosophila* homolog of vertebrate *MTG* (myeloid translocation gene) proto-oncogenes encoding nuclear scaffold proteins that form transcription repressor complexes^6–8^. In line with the high sequence similarities, pan-neuronal transgenic expression of human *MTG8* and *MTG16* significantly reduced the aggressiveness of Δ*nvy* mutants (Extended Data Fig. 2a-c), suggesting that *nvy* is a functional ortholog of these human genes. The Nvy protein harbors four Nvy Homology Regions (NHRs) that are highly conserved in mammalian MTGs^22^. We created transgenes that express Nvy proteins lacking each of the four NHRs (Extended Data Fig. 2d-e). Of the truncated *nvy* constructs, only the variant that lacked the NHR2 domain failed to rescue the Δ*nvy* phenotype (Extended Data Fig. 2f). NHR2 is required for the formation of homo-multimers of Nvy proteins (Extended Data Fig. 2g), consistent with its scaffold role in mammalian MTGs^6–8,23^. These results point to NHR2 as the key functional Nvy domain for aggression control.

## *nvy* controls social action selection

Our results indicate that *nvy* mediates the social-experience-dependent transition between aggressive states, which drives altered patterns of action selection during agonistic interactions. We next sought to characterize such changes by comparing the kinematic features of lunge-associated behavioral transitions in wild-type and Δ*nvy* flies. Although both single-reared wild-type and group-reared Δ*nvy* flies engaged in fights with a substantial number of lunges when paired with flies of the same genotype (Extended Data Fig. 3a), the Δ*nvy* mutant pairs tended to remain facing each other more often than single-reared wild-type pairs (Extended Data Fig. 3b-c). To focus our analysis on the aggressor, we hereafter isolated the attacker phenotype by pairing a “tester” fly (either single-reared wild-type or group-reared Δ*nvy*) with a group-reared wild-type “target”, which rarely lunge back (Fig. 2a). We found that the post-lunge behavioral patterns of Δ*nvy* testers were significantly distinct from those of wild-type testers. The majority of Δ*nvy* testers retained smaller facing angles (Fig. 2b-c) and distances (Extended Data Fig. 3d) to target flies after lunges, revealing that the Δ*nvy* testers remain persistently oriented after executing a lunge. To explore the link between this persistent orienting and the escalated lunging by Δ*nvy* testers, we plotted the tester’s maximum facing angle after each lunge against the latency to perform the next lunge (Fig. 2d). The Δ*nvy* testers performed more lunges at short latencies (< 2 s) without turning away (Fig. 2d, bottom), whereas the wild-type testers tended to perform lunges at intermediate intervals (2–20 s), often accompanied by large changes in facing angle (Fig. 2d, middle). When two lunges were separated by long intervals (≥ 20 s), the distributions of the maximum inter-lunge facing angle were not significantly different between the two genotypes (Fig. 2d, top). These results suggest that the *nvy* mutation primes flies to adopt a more antagonistic spatial relationship with the opponent, culminating in increased attacks. In addition to lunges, we quantified wing threat as another measure of agonistic display^24^ using the same set of movies. Intriguingly, Δ*nvy* testers exhibited shorter wing-threat duration than wild-type testers (Extended Data Fig. 3e-f), in contrast to the marked increase in lunge numbers. Given that wing threat can serve as a warning signal to the opponent^24^, this altered action choice implies that the Δ*nvy* mutant prematurely escalates the fight to direct physical attacks.

**Fig. 2.**
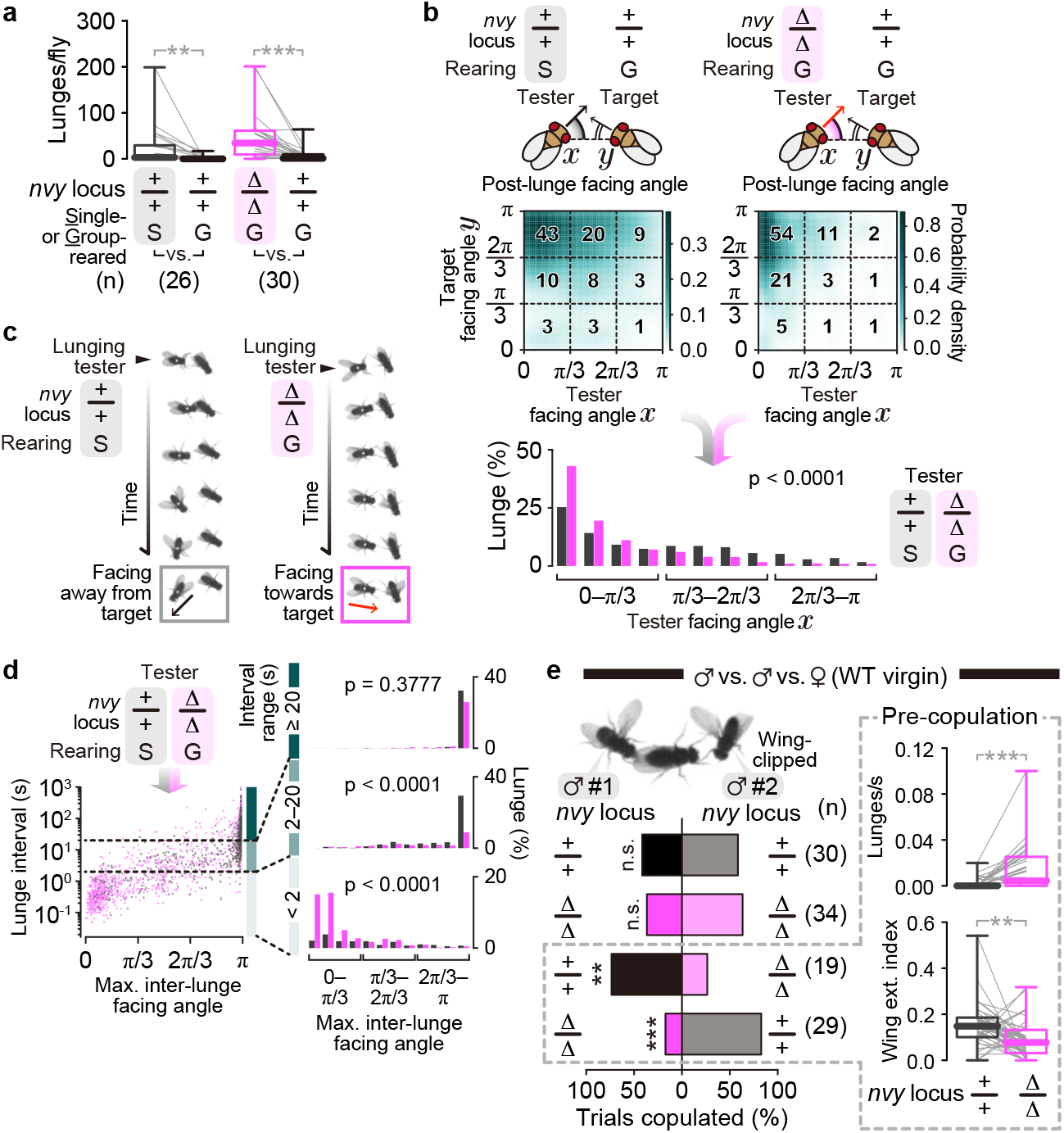
*nvy* controls agonistic action selection under various social contexts. **a**, Lunges performed by males paired for 30 min in a large chamber (see Methods). Either a Δ*nvy* male or a single-reared (“S”) wild-type male was paired with a group-reared (“G”) wild-type male. *** p < 0.0005, ** p < 0.01 (Kruskal-Wallis one-way ANOVA and post-hoc Wilcoxon signed rank test). Gray lines represent individual fly pairs. **b**, Two-dimensional heat-maps and histograms of post-lunge facing angles from movies used in **a**. Top: facing angles of tester (x axis) and target (y axis) flies 0.5 s after the end frame of each lunge are plotted as the estimated probability density. Bold numbers represent the event occurrence (%) within each of the 3 x 3 sections. Bottom: lunge occurrence (%) by the facing angles of the tester (π/12 bins). The p-value is from a Kolmogorov-Smirnov test comparing the two genotypes. **c**, Representative time-lapse images of post-lunge behaviors by wild-type (left) or Δ*nvy* (right) tester males. **d**, Relationship between post-lunge facing angles and the lunge interval. Left: scatter plot of all bouts (575 and 1,290 lunge intervals in total for the wild-type (black) and Δ*nvy* (magenta) testers, respectively) from movies used in **a**. The horizontal dashed lines indicate lunge interval ranges (short: < 2 s; intermediate: 2–20 s; long: ≥ 20 s). Right: histogram of tester’s maximum facing angles (π/12 bins) within each lunge interval range. Lunge occurrence (%) was normalized against total bouts for each genotype. The p-values are from Kolmogorov-Smirnov tests comparing the two genotypes within each lunge interval range. **e**, Competitive copulation assay using two males and one virgin female. Left: copulation success of either male #1 or #2 (wing clipped). The results of same-genotype pairs indicate that wing clipping to mark one of the males does not significantly affect their copulation tendencies. *** p < 0.0005, ** p < 0.01, n.s. p ≥ 0.05 (Fisher’s exact test). Right: lunge bouts (top) or wing extension indices (bottom) quantified in wild-type versus Δ*nvy* groups during the pre-copulation period (from the introduction of the female until either of the two males initiates copulation). *** p < 0.0005, ** p < 0.01 (Kruskal-Wallis one-way ANOVA and post-hoc Wilcoxon signed rank test). Gray lines represent individual fly groups. Numbers of pairs tested are shown in parentheses.

These results prompted us to examine the ethological roles of *nvy* in mediating aggressive action choice under a variety of social contexts. Various sensory cues from the opponent are known to affect the capacities and patterns of animal fighting behavior^4,15,25^. One such example is body size: a male fly tends to perform fewer attacks when paired against a larger male^20,26^. To test the impact of Δ*nvy* mutation on this size-dependent behavioral modulation, we generated tester males with smaller body sizes (63±8% of the target fly’s body area) by nutrition restriction and paired them with normal-sized wild-type targets. Although the small wild-type testers rarely attacked, the small Δ*nvy* flies performed lunges at the same rate as their larger targets (Extended Data Fig. 4a), eliminating the body-size effect.

We then turned to another social context in which males dramatically switch their behavioral patterns: the presence of females. Wild-type males rarely attacked females; instead, they rigorously courted them with unilateral wing extensions (Extended Data Fig. 4b), as described in earlier studies^27^. Strikingly, Δ*nvy* males performed a notable number of lunges toward females and spent less time performing unilateral wing extensions than wild-type males (Extended Data Fig. 4b). These behavioral phenotypes were reversed by pan-neuronal expression of *nvy* in Δ*nvy* males (Extended Data Fig. 4c). Despite the observed male-to-female attacks, Δ*nvy* males were as capable of copulating and forming courtship memories as wild-type males (Extended Data Fig. 4d-e). Moreover, decapitated male or female opponents provoked no lunges from Δ*nvy* males (Extended Data Fig. 4f). Therefore, the increased aggression in Δ*nvy* mutants is not likely due to sensitization to male-specific gustatory or olfactory cues, or misinterpretation of female-specific chemical cues as those of males. Compared with “one-on-one” copulation tests, a competitive assay where two males compete for one virgin female is more sensitive to subtle deficits in mating performance^28,29^. Under such conditions, the Δ*nvy* males exhibited behavioral patterns biased more towards aggression than courtship and had less copulation success than wild-type rivals (Fig. 2e). These results collectively indicate that *nvy* is necessary in flies to prevent escalated aggression, by allowing proper behavioral transitions in various ethologically relevant contexts.

## *nvy* marks aggression-suppressing neurons

To identify the neuronal mechanisms by which *nvy* suppresses aggression, we screened selected GAL4 lines to restrict *nvy* RNAi to relatively small neuronal populations. Among those tested, loss of *nvy* in neurons labeled by *Tyrosine decarboxylase 2 (Tdc2)-GAL4* increased aggression in group-reared flies most markedly (Fig. 3a-b). By contrast, *nvy* expression driven by *Tdc2-GAL4* suppressed the hyper-aggressive phenotype of Δ*nvy* (Fig. 3c). These results suggest that expression of the *nvy* gene in *Tdc2* neurons is critical for the negative regulation of aggression.

**Fig. 3.**
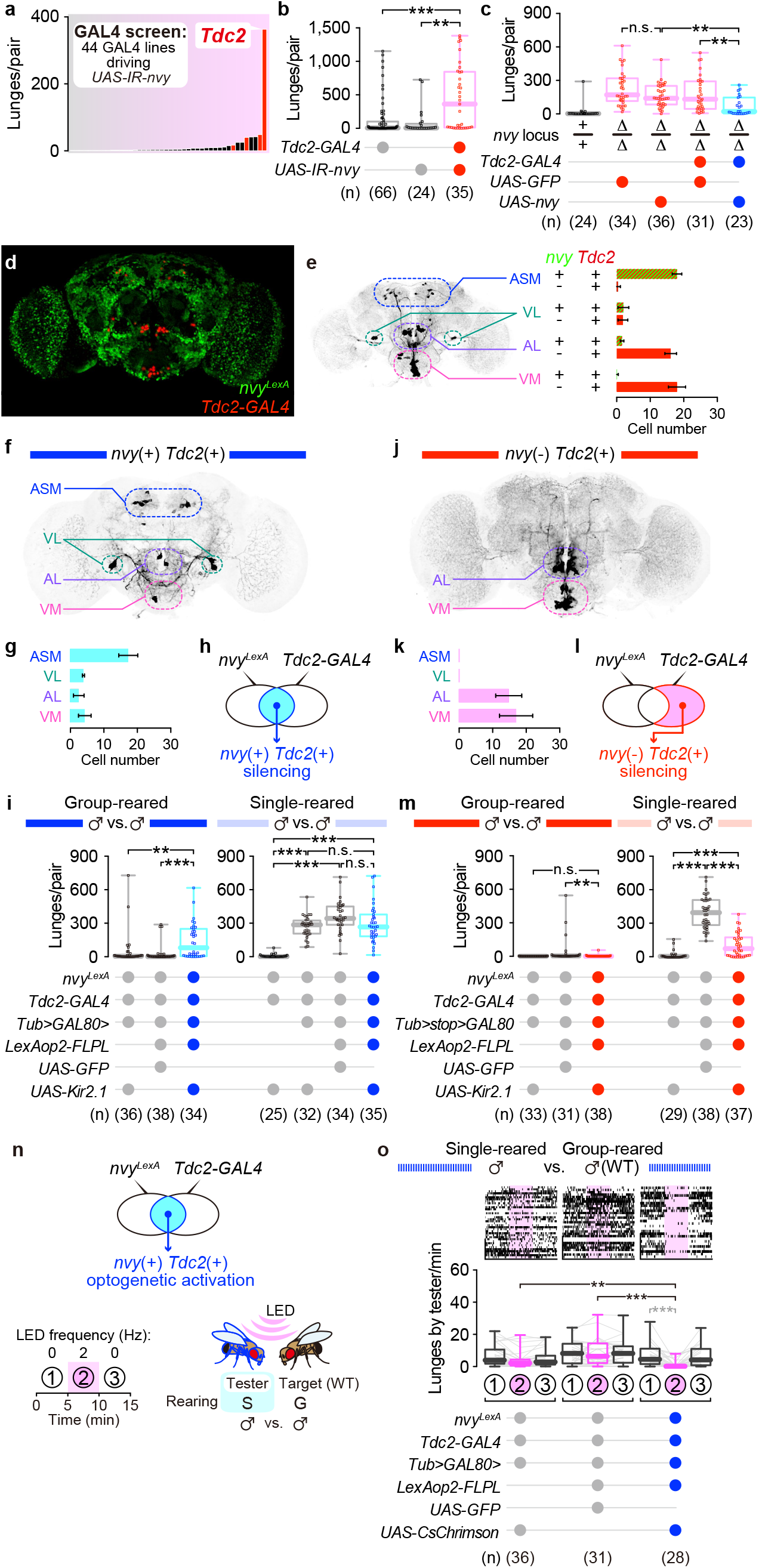
Selective manipulation of *nvy*-expressing *Tdc2* neurons suppresses aggression. **a**, GAL4 screen for neurons that suppress aggression in a *nvy*-dependent manner. Each GAL4 line expresses double-stranded RNA against *nvy*. The median numbers of lunges in 30 min by pairs of knockdown mutants (22–49 pairs for each line) are shown in the bar plot. Red bars represent 4 out of 44 GAL4 lines tested that showed a significant increase in lunges compared with two genetic controls harboring either *GAL4* or *UAS-IR*. For detailed results, see Supplementary Table S3. **b**, Increase in lunges by group-reared males in which *nvy* was knocked down by RNAi in *Tdc2* neurons. **c**, Rescue of the hyper-aggressive phenotype in Δ*nvy* males by *nvy* expression in *Tdc2* neurons. **d-e**, Co-localization of neurons labeled by *nvy^LexA^* and *Tdc2-GAL4*. Expression of nuclear localization signal (nls)-tagged GFP and tdTomato (driven by *nvy^LexA^* and *Tdc2-GAL4*, respectively), visualized by immunohistochemistry (**d**). Number of neurons co-labeled with *nvy^LexA^* and *Tdc2-GAL4* in four neuronal subtypes (ASM: anterior superior medial; VL: ventrolateral; AL: antennal lobe; VM: ventromedial), classified according to a previous anatomical study^35^ (**e**). Values represent mean ± S.D. of 8 brains. **f-m**, Selective silencing of *nvy*-positive or *nvy*-negative *Tdc2* neurons in males. Expression of GFP in *nvy*-positive (**f**) or *nvy*-negative (**j**) *Tdc2* neurons visualized by immunohistochemistry, and cell counts for each subpopulation (mean ± S.D. of 10 brains for each genotype; **g**, **k**). Genetic access to each subpopulation was achieved by either intersection (**h**) or subtraction (**l**) of *Tdc2-GAL4* and *nvy^LexA^* (see Methods). Altered aggression after Kir2.1-mediated silencing of either *nvy*-positive or *nvy*-negative *Tdc2* neurons is shown in box plots (**i, m**). For each genotype (indicated below the box plots), lunges were quantified in group-reared (left) or single-reared (right) male pairs. **n**, Optogenetic stimulation paradigm. Each experiment consisted of three time windows: the first 5 min without stimulation (“1”), the next 5 min with stimulation (“2”), and the last 5 min without stimulation (“3”). Single-reared tester males were paired with group-reared wild-type target males. **o**, Reduced aggression in socially naïve males by optogenetic stimulation of the *nvy*-positive *Tdc2* neurons. Raster plots (top) and box plots (bottom) of lunges performed by the tester males are shown. In black: *** p < 0.0005, ** p < 0.005, n.s. p ≥ 0.05 (Kruskal-Wallis one-way ANOVA and post-hoc Mann-Whitney U-test with Bonferroni correction). In gray: *** p < 0.0005 (Kruskal-Wallis one-way ANOVA and post-hoc Wilcoxon signed rank test).

*Tdc2* encodes the biosynthetic enzyme for octopamine/tyramine (OA/TA)^30^, the invertebrate counterparts of norepinephrine/epinephrine. In flies, OA/TA regulate a wide variety of behaviors including aggression^31,32^. Since OA itself and octopaminergic neurons have been reported to promote fly aggression^12,20,31–33^, identification of *Tdc2* neurons as the site of *nvy*-dependent *suppression* of aggression intrigued us. Consistent with a previous report^20^, expressing an inward rectifying potassium channel *Kir2.1*^34^ (which induces neuronal hyperpolarization) in all *Tdc2* neurons suppressed the basal aggressiveness of single-reared flies (Extended Data Fig. 5a). To examine whether *nvy* is expressed in all *Tdc2* neurons or just a subset, we gained genetic access to the *nvy*-expressing cells by creating a CRISPR/Cas9-mediated knock-in allele of the *nvy* locus that expresses the bacterial transcription factor *LexA* in place of *nvy* (*nvy^LexA^*; Extended Data Fig. 5b-c). This knock-in allele is null for *nvy*, as heteroallelic combination of *nvy^LexA^* and Δ*nvy* reduced Nvy expression to an undetectable level and resulted in hyper-aggressiveness, similarly to homozygous Δ*nvy* (Extended Data Fig. 5d-e). Intersection of *Tdc2-GAL4* and *nvy^LexA^* labeled a specific, *nvy*-expressing subset of *Tdc2* neurons^35^ (Fig. 3d-e).

We first suppressed the neuronal activity of either the *nvy^LexA^*-positive or -negative *Tdc2* populations by selectively expressing *Kir2.1*. Group-reared males that normally have low basal aggressiveness showed a marked increase in lunges when the *nvy^LexA^*-positive *Tdc2* population was silenced (Fig. 3f-i). This genetic intersection therefore labels aggression-suppressing neurons. By contrast, the high intensity of basal aggression in single-reared males was not affected by the same manipulation (Fig. 3i), similar to the Δ*nvy* mutant phenotype (Fig. 1e). Silencing of *nvy^LexA^*-negative *Tdc2* cells had opposing effects; group-reared males remained non-aggressive whereas single-reared males performed fewer lunges (Fig. 3j-m). These data reveal that *Tdc2* neurons contain a *nvy*-expressing population required to suppress aggression and a non-*nvy*-expressing population that can promote aggression. This functional heterogeneity within aminergic neurons provides flexibility in control of aggression according to social experience. Our findings parallel recent vertebrate studies reporting lateral habenula subpopulations that respond in opposite directions during aggressive encounters^36,37^.

The above chronic silencing results encouraged us to probe the aggression-suppressing role of *Tdc2* neurons in socially naïve animals using optogenetics, which allows for greater temporal control of neural activity. Single-reared testers that express the channelrhodopsin CsChrimson^38^ specifically in the *nvy^LexA^*-positive *Tdc2* population were photostimulated by a red light-emitting diode (Fig. 3n). We found that the basal aggressiveness of tester males during the stimulation period was significantly lower than genetic controls (Fig. 3o and Extended Data Fig. 6a-b). General locomotion, “time orienting” (the length of time a tester fly is in close proximity to and likely to interact with a target fly^39,40^), and male-to-female courtship were largely unaffected by stimulation (Extended Data Fig. 6c-h). Thus, activation of the *nvy^LexA^*-positive *Tdc2* population specifically blocks the execution of lunges without affecting other social behaviors. Our data collectively indicate that the previously uncharacterized *nvy*-expressing *Tdc2* subpopulation serves as a neuronal switch that controls social-experience-dependent changes in aggressiveness.

Similar to males, female flies also alter their aggressiveness according to their social experience^41^. To investigate whether our findings on the role of *nvy* are common to both sexes, we applied the above genetic and neuronal manipulations to female flies. Female aggressiveness was assessed by quantifying headbutts, a female-type aggressive action^42^. The Δ*nvy* mutation increased headbutts in group-reared females (Fig. 4a), and this increase was suppressed by transgenic *nvy* expression (Fig. 4b and Extended Data Fig. 7a). As with males, the elevated basal aggressiveness of single-reared wild-type females was reduced by overexpression of *nvy* (Fig. 4c). Moreover, both silencing (Fig. 4d and Extended Data Fig. 7b-c) and optogenetic activation (Fig. 4e) of the *nvy*-expressing *Tdc2* population in females had similar effects on aggression as in males. The observed commonality in behavioral responses in males and females is consistent with the similar morphologies of *Tdc2* neurons in the central brain of both sexes (Extended Data Fig. 7d-e). Although sexually dimorphic neurons under control of the sex-determining genes *fruitless* and *doublesex* have been widely studied with respect to fly social behaviors^9,11,13,43–47^, it is relatively rare that neurons specified by the same genetic agent affect aggression in both sexes^10^. A recent study in mice demonstrated that a subpopulation of neurons in the ventromedial hypothalamus, previously known as a region essential for male aggression^48^, controls aggression in females as well^49^. Our work provides an entry point to elucidate unexplored common mechanisms underlying sexually dimorphic behaviors.

**Fig. 4.**
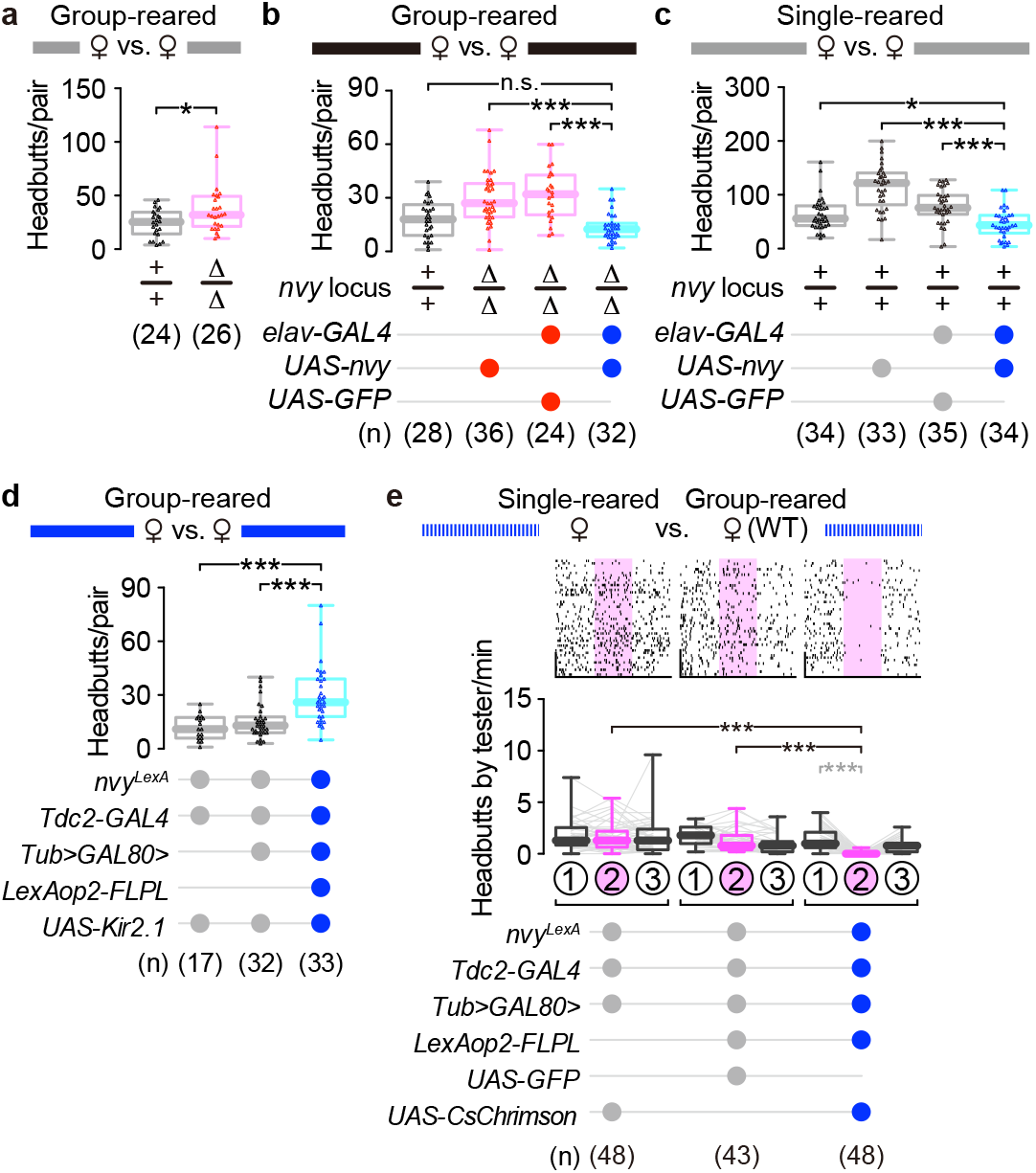
The *nvy* gene and *nvy*-positive *Tdc2* neurons suppress aggression in females. **a**, The Δ*nvy* mutation increases aggression in socially experienced females. Headbutts performed in 30 min by group-reared virgin female pairs. **b**, Rescue of the hyper-aggressive phenotype in Δ*nvy* females by the pan-neuronal *nvy* expression. **c**, Reduced aggression in socially naïve females by the pan-neuronal *nvy* overexpression. **d**, Increased aggression in socially experienced females after Kir2.1-mediated silencing of *nvy*-positive *Tdc2* neurons. **e**, Reduced aggression in socially naïve females by optogenetic stimulation of *nvy*-positive *Tdc2* neurons. Single-reared tester females with the indicated genotypes were paired with group-reared wild-type target females. The optogenetic stimulation paradigm was same as in **Fig. 3n**. Raster plots (top) and box plots (bottom) of headbutts performed by the tester females are shown. In black: *** p < 0.0005, * p < 0.05, n.s. p ≥ 0.05 (Kruskal-Wallis one-way ANOVA and post-hoc Mann-Whitney U-test with Bonferroni correction). In gray: *** p < 0.0005 (Kruskal-Wallis one-way ANOVA and post-hoc Wilcoxon signed rank test).

## Aggression-regulating genes downstream of *nvy*

We next explored the molecular mechanism through which *nvy* acts within in *Tdc2* neurons. As demonstrated above, the aggression-suppressing effect of Nvy requires NHR2, the conserved domain critical for transcriptional regulatory functions of vertebrate MTG proteins^6–8^. We therefore focused on the transcriptional activity of *Tdc2* neurons, comparing the gene expression profiles of wild-type and Δ*nvy Tdc2* neurons using single-cell RNA sequencing. The number of cells labeled by *Tdc2-GAL4* was comparable in wild-type and Δ*nvy* brains (Extended Data Fig. 8a-b). These *Tdc2* cells labeled by membrane-bound GFP derived from either wild-type or Δ*nvy* background flies were harvested from brain homogenates using a fluorescence-activated cell sorter (Extended Data Fig. 8c). Hierarchical clustering analysis^50^ on the data collected from 171 *Tdc2* cells (82 from wild-type and 89 from Δ*nvy*; Extended Data Fig. 8d) revealed 6 clusters with distinct expression patterns (Fig. 5a and Extended Data Fig. 8e-i). Among them, cluster #5 was enriched in cells expressing *nvy* at relatively high levels (Extended Data Fig. 8i-k; note that the *nvy* mRNA is detectable in homozygous Δ*nvy* as the mutation removes only the 1st exon). We performed differentially expressed gene (DEG) analysis on this cluster. Comparison of wild-type and Δ*nvy* cells within cluster #5 identified 25 down-regulated and 12 up-regulated genes with fold changes greater than 10 (Fig. 5b), whereas no such DEGs were detected by analyzing all *Tdc2* cells as a whole (Extended Data Fig. 8l). These data suggest that *nvy* affects gene expression in a cell-type-specific manner.

**Fig. 5.**
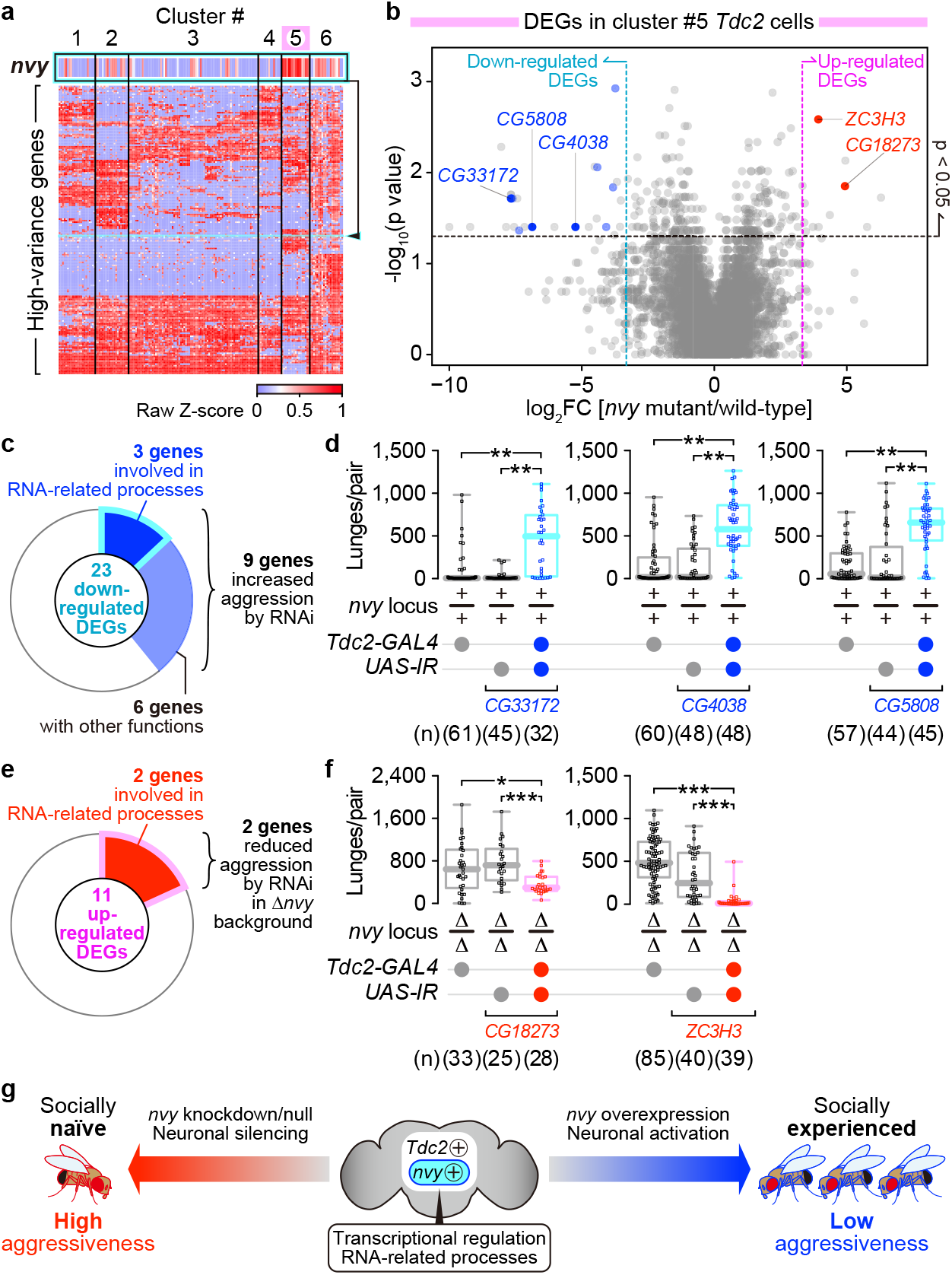
*nvy* functions in *Tdc2* neurons to control aggression *via* transcriptional modulation. **a**, Hierarchical iterative clustering analysis of *Tdc2* cells based on gene expression values from single-cell RNA sequencing. Of the top 200 high-variance genes listed in the vertical direction, the row showing the expression pattern of *nvy* is magnified. Six resulting clusters, including the one most enriched with *nvy*-expressing cells (cluster #5), are labeled at the top. **b**, Volcano plot of DEGs in cluster #5 cells. Dots are plotted according to the fold change (FC) and the p-value (by Mann-Whitney U-test) of each gene when the Δ*nvy* mutant cells were compared against the wild-type cells. Pale-colored dots represent genes that pass the Benjamini-Hochberg FDR test (blue: down-regulated at FC < 0.1; red: up-regulated at FC > 10); dark-colored dots correspond to genes that showed behavioral phenotypes in the following RNAi experiments **(c-f). c**, Down-regulated DEGs found in cluster #5 that showed increased aggression with RNAi in the wild-type *nvy* background. **d**, Increased aggression by *Tdc2-GAL4-*driven RNAi of three down-regulated DEGs with predicted roles in RNA-related processes. ** p < 0.01, * p < 0.05 (Kruskal-Wallis one-way ANOVA and post-hoc Mann-Whitney U-test with Bonferroni correction). **e**, Up-regulated DEGs found in cluster #5 that showed decreased aggression with RNAi in the Δ*nvy* background. **f**, Reduced aggression in the Δ*nvy* mutants following the *Tdc2-GAL4-*driven RNAi of two up-regulated DEGs predicted to be involved in RNA-related processes. *** p < 0.0005, * p < 0.05 (Kruskal-Wallis one-way ANOVA and post-hoc Mann-Whitney U-test with Bonferroni correction). **g**, A schematic summary of the modulation of social experience-dependent aggression through manipulation of *nvy* and *nvy*-expressing *Tdc2* neurons.

If *nvy* controls aggression through transcriptional regulation, DEGs found in the *nvy*-expressing *Tdc2* neurons may contain effector genes necessary for appropriate modulation of aggression. Supporting this idea, *Tdc2-GAL4*-driven RNAi targeting 9 down-regulated protein-coding genes found in cluster #5 significantly increased lunge numbers in wild-type flies under group-rearing conditions (Fig. 5c-d and Extended Data Fig. 9a-c), phenocopying the *nvy* RNAi. Knocking down 2 up-regulated genes reduced the aggressiveness of Δ*nvy* mutants (Fig. 5e-f and Extended Data Fig. 9d-f), suggesting that these genes normally act downstream of *nvy* to suppress aggression. It is noteworthy that 5 DEGs that showed behavioral phenotypes have proposed roles in RNA-related processes (Extended Data Fig. 9g). The fact that genes controlled by *nvy* can both promote and suppress aggression suggests that *nvy* serves as a molecular hub that coordinates changes in the level of aggression through a specific subpopulation of *Tdc2* neurons.

Although genetic predispositions influence basal levels of aggression, animals from relatively homogeneous genetic backgrounds can exhibit a wide range of aggressiveness^1,51^. This variability may be the result of social experiences that are unique to each individual. Our findings underscore the importance of an active suppression mechanism for properly adjusting the intensity of aggressive behavior according to social experience (in this case, group rearing), and for preventing excess aggression that is maladaptive in certain social contexts (Fig. 5g). The identification of *nvy* and *nvy*-expressing *Tdc2* neurons as novel genetic and neuronal “brakes” for aggression in a sex-invariant manner advances our understanding of the elaborate neural mechanisms that allow animals to flexibly modulate aggressive behaviors in accordance with their social environments.

---

**Extended Data Fig. 1.**
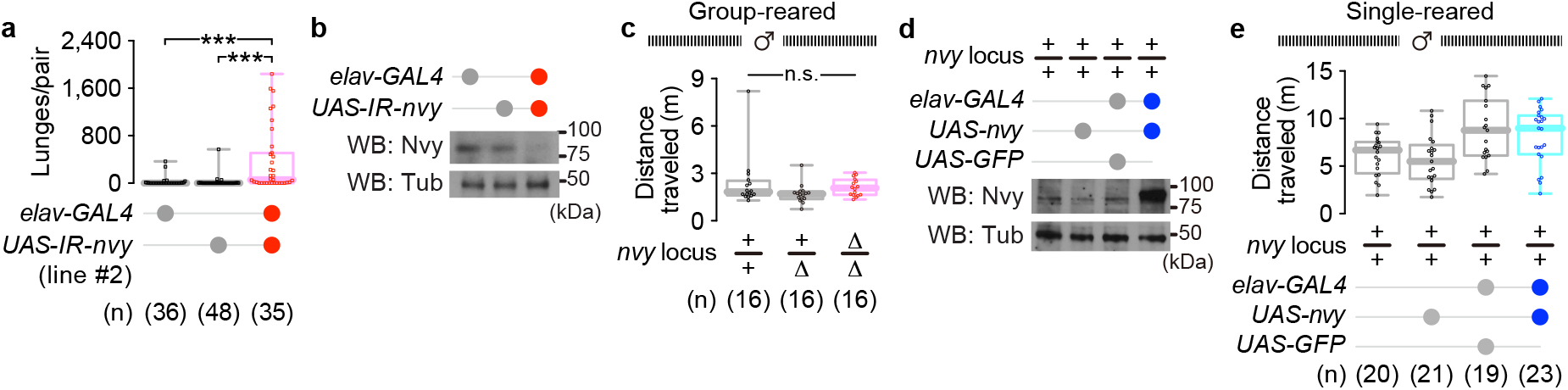
Additional behavioral characterization of *nvy* mutants. **a**, Increased lunges performed by males following pan-neuronal knockdown of *nvy* by another *UAS-IR* strain (JF03349). *** p < 0.0005 (Kruskal-Wallis one-way ANOVA and post-hoc Mann-Whitney U-test with Bonferroni correction). **b**, Reduced expression of Nvy protein in fly heads following the pan-neuronal knockdown of *nvy*, verified by Western blot. α-Tubulin (Tub) was used as an internal control. **c**, Locomotor activity of Δ*nvy* males was not affected. Group-reared males of the indicated genotypes were introduced individually into the chamber and the distance traveled in 30 min was measured. n.s. p ≥ 0.05 (Kruskal-Wallis one-way ANOVA). **d**, Expression of Nvy protein in fly heads following pan-neuronal overexpression of *nvy*, verified by Western blot. **e**, Locomotor activity of flies following pan-neuronal overexpression of *nvy* was comparable to genetic controls. Single-reared males were introduced individually into the chamber and the distance traveled in 30 min was measured.

**Extended Data Fig. 2.**
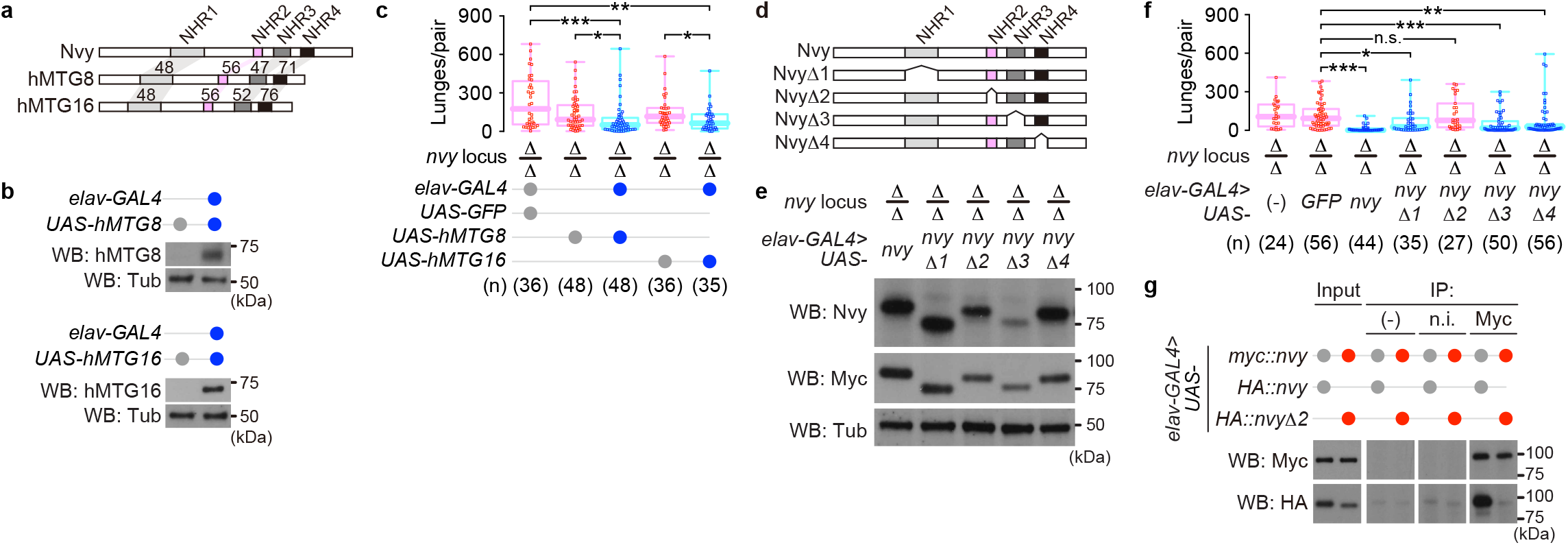
Additional biochemical and behavioral data for human MTGs and truncated versions of Nvy. **a**, Gene structures of human *MTGs* with amino acid sequence identities (%) against *nvy* within each NHR domain. **b**, Pan-neuronal expression of human *MTG* genes by *elav-GAL4*, verified in fly head extracts by Western blot. **c**, Rescue of the Δ*nvy* phenotype by pan-neuronal expression of human *MTGs* under control of *UAS*. *** p < 0.001, ** p < 0.01, * p < 0.05 (Kruskal-Wallis one-way ANOVA and post-hoc Mann-Whitney U-test with Bonferroni correction). **d**, Schematic of the truncated *UAS-nvy* constructs lacking each NHR domain. **e**, Pan-neuronal expression of mutated *nvy* transgenes lacking one of the NHR1–4 domains in the Δ*nvy* background. All *UAS-nvy* constructs contain 3xMyc tags at the N-terminus. α-Tubulin (Tub) was used as an internal control. **f**, Rescue of the Δ*nvy* phenotype by pan-neuronal expression of truncated *UAS-nvy* constructs. *** p < 0.0005, ** p < 0.005, * p < 0.05, n.s. p ≥ 0.05 (Kruskal-Wallis one-way ANOVA and post-hoc Mann-Whitney U-test with Bonferroni correction). **g**, Homo-multimer formation of Nvy protein mediated by the NHR2 domain. Myc-tagged Nvy was co-immunoprecipitated with either HA-tagged intact Nvy or the mutated version lacking NHR2 (*nvy*Δ*2*). Input: 7.5% of lysate used for the precipitation. IP, (-): samples precipitated with no antibody. IP, n.i.: samples precipitated with normal IgG. IP, Myc: samples precipitated with an anti-Myc antibody.

**Extended Data Fig. 3.**
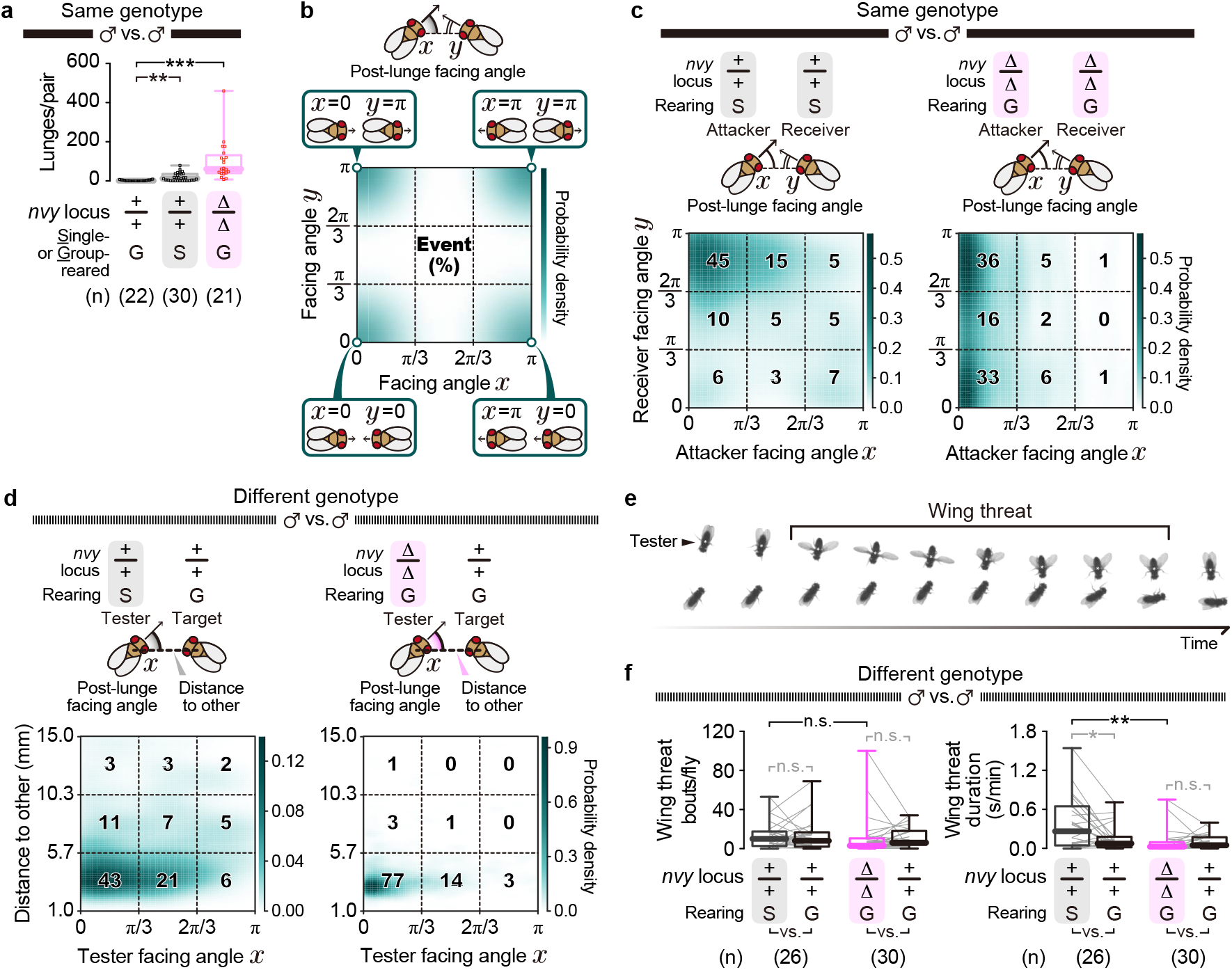
Additional pairing conditions, kinematic features, and behavior types analyzed in the agonistic action selection experiments. **a**, Lunges performed by males in a large chamber during the 30-min recording period. Genotypes and rearing conditions are shown below the plots. *** p < 0.0005, ** p < 0.005 (Kruskal-Wallis one-way ANOVA and post-hoc Mann-Whitney U-test with Bonferroni correction). **b**, Schematic of the two-dimensional heat-map used for display post-lunge facing angles. **c**, Two-dimensional probability-density plots of post-lunge facing angles in same-genotype pairs, from the same movies used in **a**. Facing angles of the fly either performing (“attacker”, x axis) or receiving a lunge (“receiver”, y axis) 0.5 s after the end frame of each lunge are plotted as the estimated probability density (scale to the right). Bold numbers represent the event occurrence (%) within each 3 x 3 section. **d**, Two-dimensional plots of the post-lunge facing angle of the tester flies (x axis) and the distance between the two flies (y axis) 0.5 s after each lunge, plotted as the estimated probability density (scale to the right). Original movies used were the same as those in **Fig. 2a-e**. Bold numbers represent the event occurrence (%) within each 3 x 3 section. **e**, Representative time-lapse frames of a tester male displaying wing threat. **f**, Bout numbers (left) and duration (right) of wing threat displayed by testers paired with group-reared wild-type targets. Original movies used were the same as those in **Fig. 2a-e**. In black: ** p < 0.005, n.s. p ≥ 0.05 (Kruskal-Wallis one-way ANOVA and post-hoc Mann-Whitney U-test). In gray: * p < 0.05, n.s. p ≥ 0.05 (Kruskal-Wallis one-way ANOVA and post-hoc Wilcoxon signed rank test).

**Extended Data Fig. 4.**
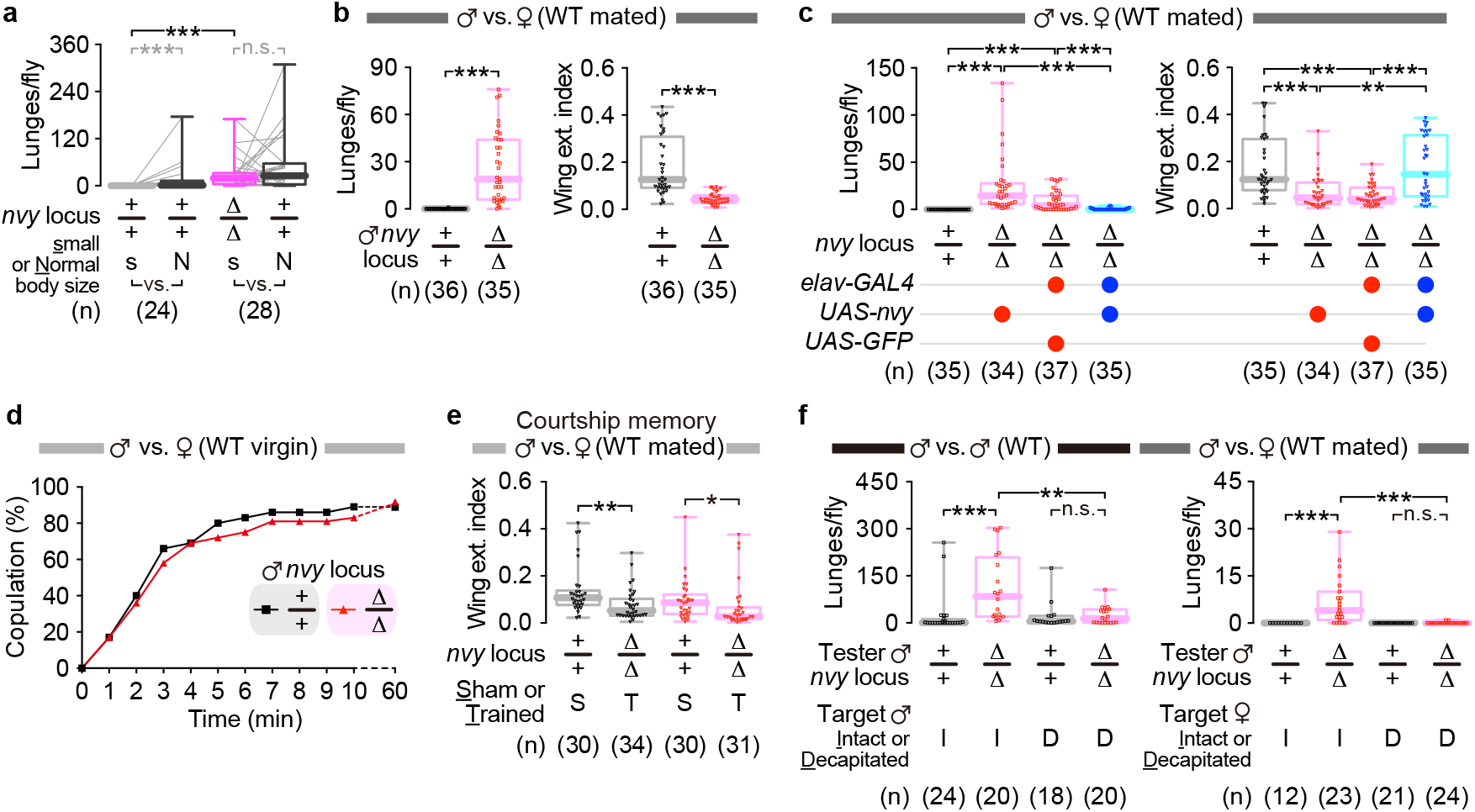
Behavior phenotypes of Δ*nvy* males under various social contexts. **a**, Lunges performed in 30 min by pairs of small-(“s”) and normal-(“N”) sized males. In black: *** p < 0.0005 (Kruskal-Wallis one-way ANOVA and post-hoc Mann-Whitney U-test). In gray: *** p < 0.0005, n.s. p ≥ 0.05 (Kruskal-Wallis one-way ANOVA and post-hoc Wilcoxon signed rank test). **b**, Aggression and courtship behaviors performed by males against wild-type mated females. Lunge numbers (left) and wing extension indices (right; the relative duration of time spent performing wing extensions) measured over 1 h. *** p < 0.0005 (Kruskal-Wallis one-way ANOVA and post-hoc Mann-Whitney U-test). **c**, Rescue of the male-to-female behavioral phenotypes in Δ*nvy* by pan-neuronal expression of *nvy*. Lunge numbers (left) and wing extension indices (right) measured over 1 h. Genotypes of male testers are indicated below the plots. *** p < 0.0005, ** p < 0.005 (Kruskal-Wallis one-way ANOVA and post-hoc Mann-Whitney U-test with Bonferroni correction). **d**, Cumulative copulation rates of males paired with wild-type virgin females (n = 24). **e**, Courtship memory assay. Wing extension indices of males during the test session. Males were previously either trained with mated females (“T”) or sham-trained (“S”). ** p < 0.005, * p < 0.05 (Kruskal-Wallis one-way ANOVA and post-hoc Mann-Whitney U-test with Bonferroni correction). **f**, Aggression by Δ*nvy* males requires behavioral feedback from the opponent. Lunges in 30 min by either wild-type or Δ*nvy* tester males, toward intact (“I”) or decapitated (“D”) wild-type target males (left) or mated females (right). *** p < 0.0005, ** p < 0.005, n.s. p ≥ 0.05 (Kruskal-Wallis one-way ANOVA and post-hoc Mann-Whitney U-test with Bonferroni correction).

**Extended Data Fig. 5.**
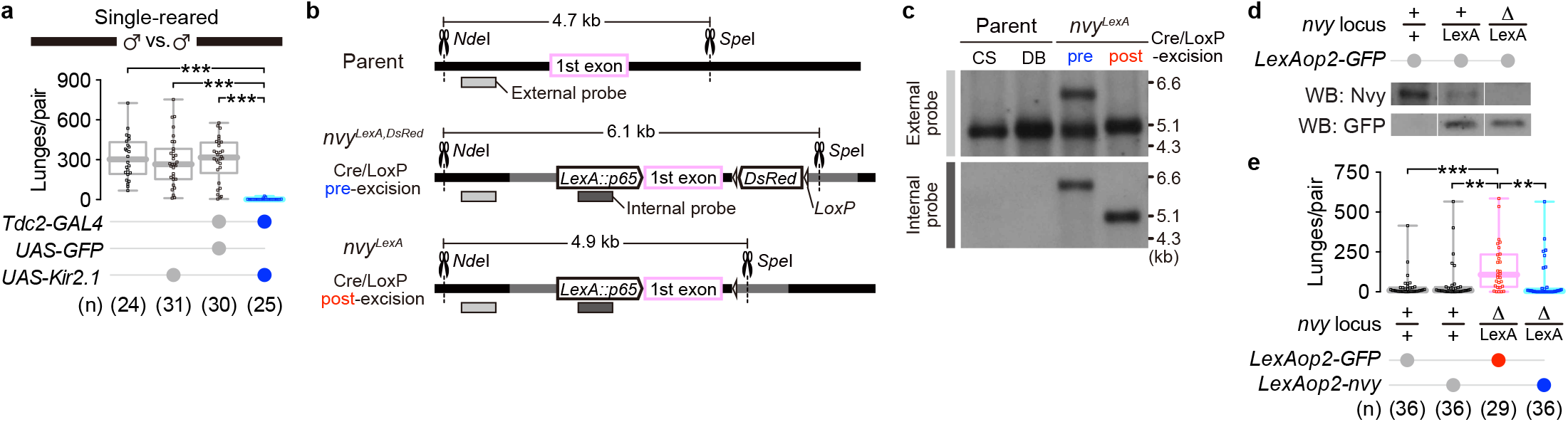
Generation of the *nvy^LexA^* knock-in lines and additional behavioral data for the *Tdc2* silencing experiments. **a**, Reduced aggression in socially naïve males by Kir2.1-mediated silencing of all *Tdc2* cells. Lunges performed by single-reared male pairs. *** p < 0.0005 (Kruskal-Wallis one-way ANOVA and post-hoc Mann-Whitney U-test with Bonferroni correction). **b**, Genome schematics of the *nvy^LexA^* knock-in alleles. The *nvy* locus of the parental line (top) was targeted by CRISPR/Cas9-mediated cleavage, leading to homologous recombination with the plasmid harboring the coding sequences of *LexA::p65* and the eye-specific genetic marker *3XP3-DsRed* (middle). After backcrossing with the wild-type strain, the *DsRed* marker flanked by *LoxP* was excised by crossing with an *hs-Cre* line (bottom). Predicted distances between the recognition sites of two restriction enzymes, *Nde*I and *Spe*I, are shown for each genotype. For the following Southern blot analysis, one region outside the *nvy* exon and another region inside the *LexA::p65* coding sequence were targeted by “external” and “internal” probes, respectively. **c**, Southern blot analysis of the parental and *nvy^LexA^* knock-in lines. *Nde*I/*Spe*I-digested genomic DNA from each line was hybridized with either the external (top) or internal (bottom) probe. For parental lines, the wild-type Canton-S (CS) used for backcrossing and the “double-balancer” (DB: *w*; *Bl*/*CyO*; *TM2*/*TM6B*) used to establish the knock-in lines are shown. Note that the *nvy^LexA^* knock-in lines were maintained with the second chromosome balancer *CyO* derived from the parental DB line. **d**, Western blot analysis of Nvy protein extracted from the *nvy^LexA^* fly heads. **e**, Hyperaggressive phenotype induced by trans-heterozygosity of *nvy^LexA^* and Δ*nvy*, and its rescue by *nvy* expression. *** p < 0.0005, ** p < 0.005 (Kruskal-Wallis one-way ANOVA and post-hoc Mann-Whitney U-test with Bonferroni correction).

**Extended Data Fig. 6.**
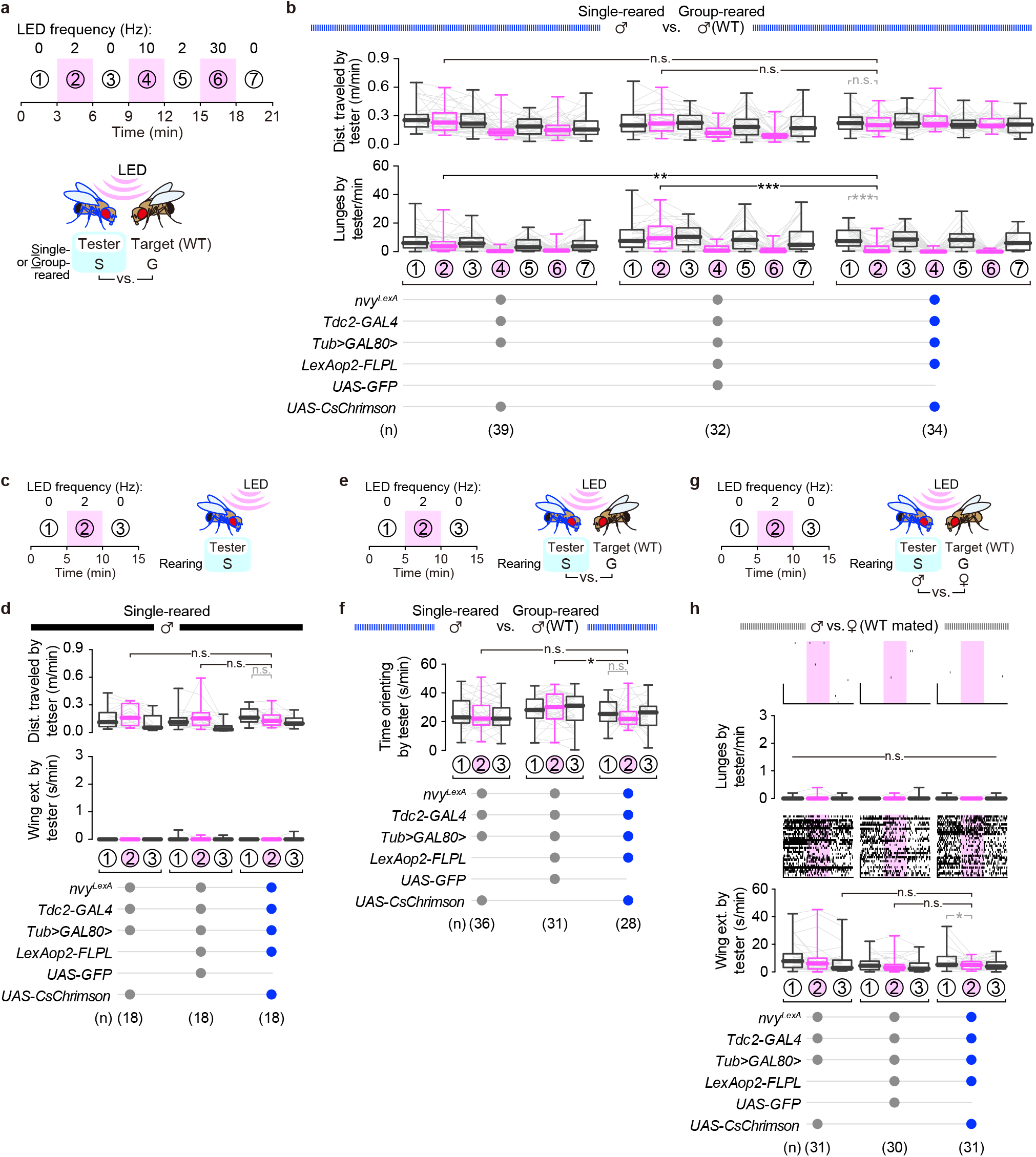
Additional behavioral data obtained during optogenetic stimulation of *nvy*-positive *Tdc2* neurons. **a-b**, Optogenetic stimulation of *nvy*-positive *Tdc2* neurons at various LED frequencies. The stimulation was performed at 2, 10, and 30 Hz for 3 min, each separated by a 3-min interval (**a**). Distance traveled (**b**; top) and lunges (**b**; bottom) performed by tester males during each 3-min time window are shown in box plots. In black: *** p < 0.0005, ** p < 0.01, n.s. p ≥ 0.05 (Kruskal-Wallis one-way ANOVA and post-hoc Mann-Whitney U-test with Bonferroni correction). In gray: *** p < 0.0005, n.s. p ≥ 0.05 (Kruskal-Wallis one-way ANOVA and post-hoc Wilcoxon signed rank test). **c-d**, Optogenetic stimulation of *nvy*-positive *Tdc2* neurons in solitary testers. Stimulation was performed at 2 Hz for 5 min in the absence of a target fly (**c**). Distance traveled (**d**; top) and wing extensions (**d**; bottom) performed by tester males during each 5-min time window. In black: n.s. p ≥ 0.05 (Kruskal-Wallis one-way ANOVA and post-hoc Mann-Whitney U-test with Bonferroni correction). In gray: n.s. p ≥ 0.05 (Kruskal-Wallis one-way ANOVA and post-hoc Wilcoxon signed rank test). **e-f**, Orienting toward target flies during optogenetic stimulation of *nvy*-positive *Tdc2* neurons (**e**). The original movies used in **Fig. 3o** were reanalyzed. Time spent by tester males orienting towards the target males during each 5-min window (**f**). In black: * p < 0.05, n.s. p ≥ 0.05 (Kruskal-Wallis one-way ANOVA and post-hoc Mann-Whitney U-test with Bonferroni correction). In gray: n.s. p ≥ 0.05 (Kruskal-Wallis one-way ANOVA and post-hoc Wilcoxon signed rank test). **g-h**, Male-to-female lunges and wing extensions during optogenetic stimulation of *nvy*-positive *Tdc2* neurons. Male testers were paired with wild-type mated females, and the stimulation was performed at 2 Hz for 5 min (**g**). Lunges (**h**, top) and wing extensions (**h**, bottom) performed by target males during each 5-min window. The pink area within each raster plot indicates the stimulation period (time window “2” in **g**). In black: n.s. p ≥ 0.05 (Kruskal-Wallis one-way ANOVA). In gray: * p < 0.05 (Kruskal-Wallis one-way ANOVA and post-hoc Wilcoxon signed rank test).

**Extended Data Fig. 7.**
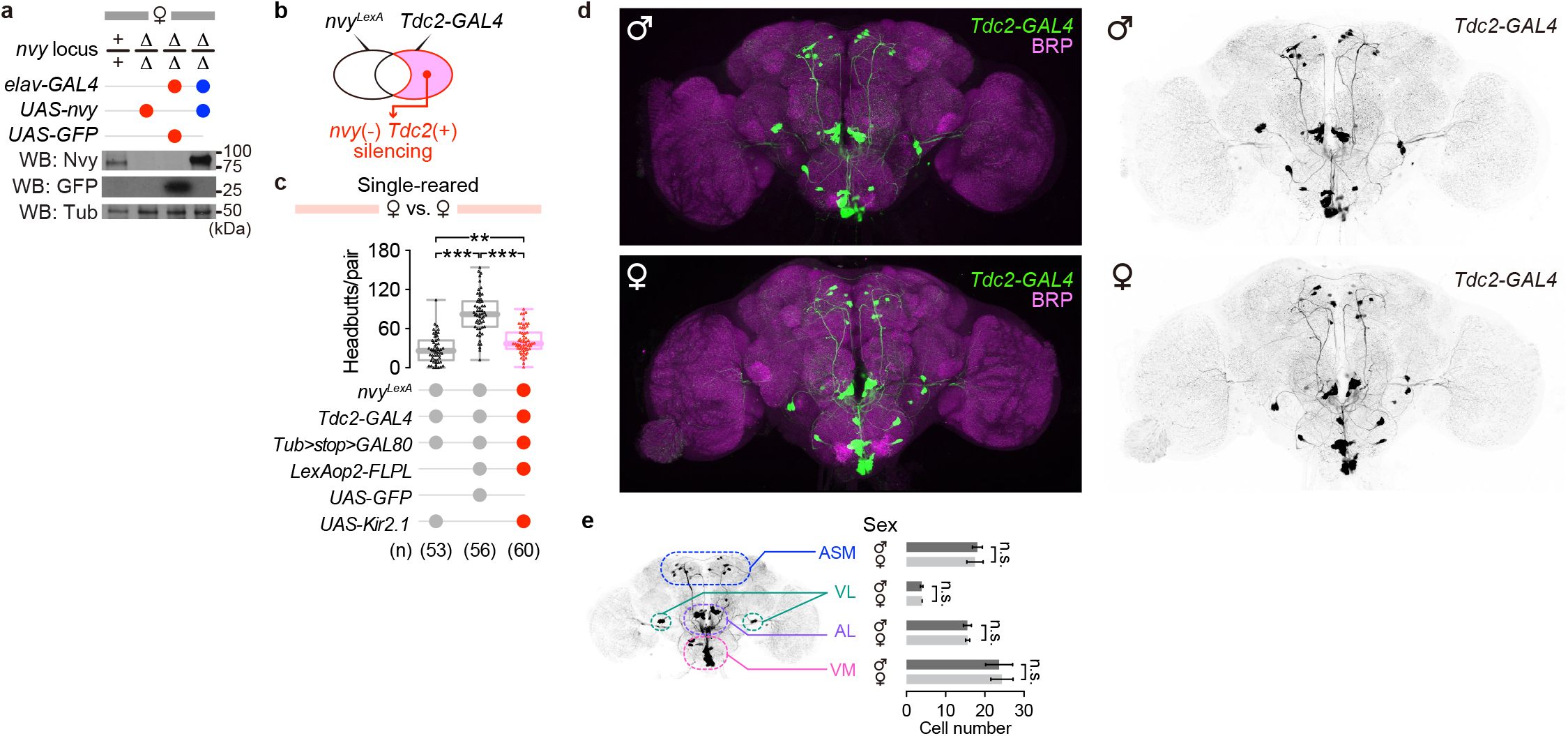
Additional expression and behavioral data from experiments with females. **a**, Pan-neuronal expression of Nvy in Δ*nvy* background females verified by Western blot. α-Tubulin (Tub) was detected as an internal control. **b-c**, Selective silencing of *nvy*-negative *Tdc2* neurons in females. Genetic access to the *nvy*-negative subpopulation was achieved by subtraction between *Tdc2-GAL4* and *nvy^LexA^* (**b**; see Methods). Headbutts in socially naïve females are shown in box plots (**c**). *** p < 0.0005, ** p < 0.005 (Kruskal-Wallis one-way ANOVA and post-hoc Mann-Whitney U-test with Bonferroni correction). **d,** Neuronal morphology of *Tdc2* neurons in male and female brains. Left: GFP expressed under the control of *Tdc2-GAL4*, along with the neuropil marker Bruchpilot (BRP), were visualized by immunohistochemistry using male (top) or female (bottom) brains. Right: same images as left with GFP signals visualized in gray scale. **e**, Cell counts of *Tdc2* neurons in males and females. Subtypes of *Tdc2-GAL4* neurons were classified according to a previous anatomical study^35^. Values represent mean ± S.D. of 9 (male) or 8 (female) brains. n.s. p ≥ 0.05 (unpaired t-test).

**Extended Data Fig. 8.**
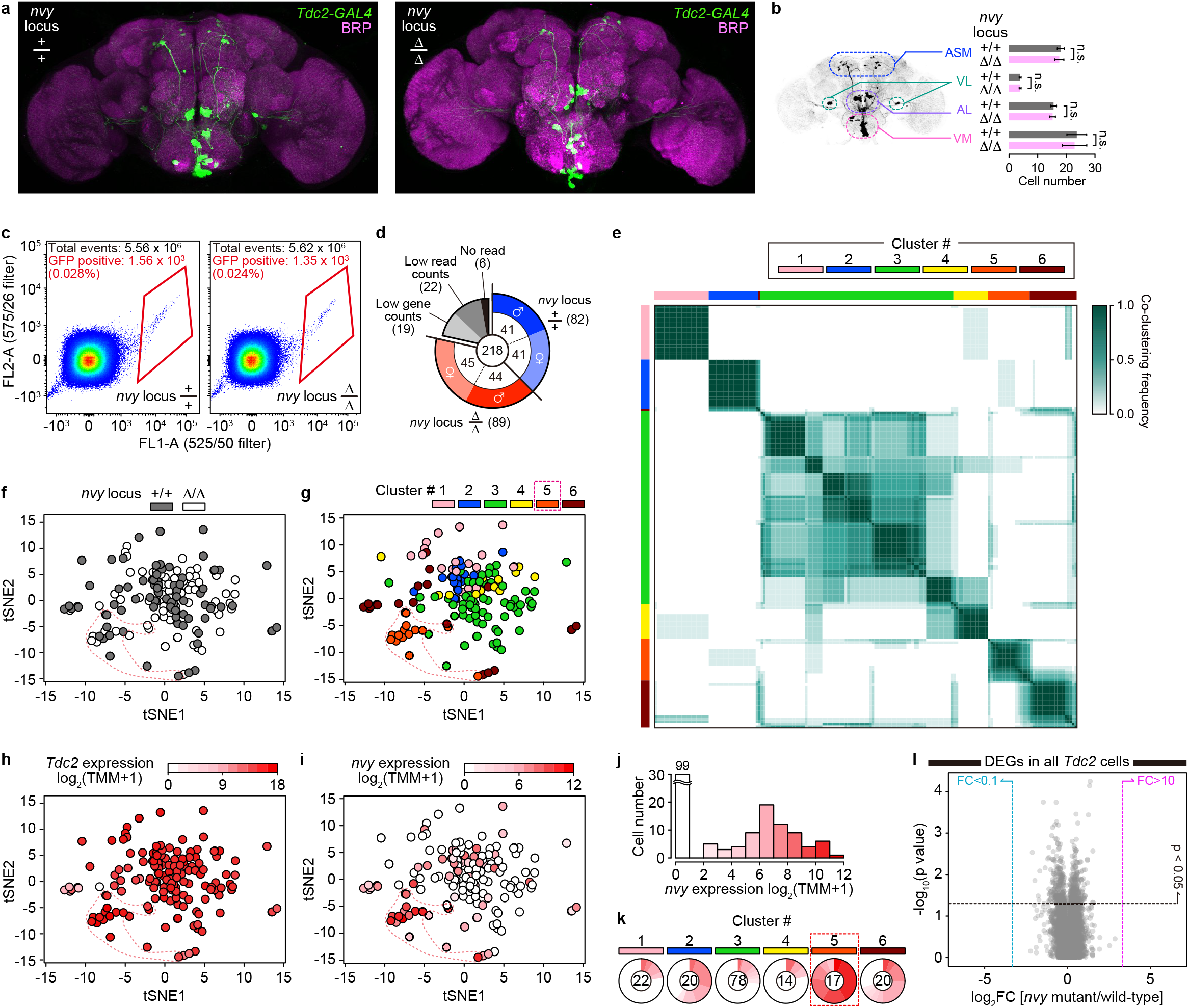
Additional analyses of cell clusters and DEGs from single-cell RNA-sequencing of *Tdc2* neurons. **a**, Neuronal morphology of *Tdc2* neurons in the Δ*nvy* brain. GFP expressed under the control of *Tdc2-GAL4*, along with the neuropil marker BRP, were visualized by immunohistochemistry in brains from *nvy* locus wild-type (left) or Δ*nvy* (right) males. **b**, Cell counts of *Tdc2* neurons in the Δ*nvy* brain. Subtypes of *Tdc2-GAL4* neurons were classified according to a previous anatomical study^35^. Note that the values for the wild-type *nvy* locus are re-plotted from Extended Data Fig. 7e. Values represent mean ± S.D. of 9 brains. n.s. p ≥ 0.05 (unpaired t-test). **c**, FACS results for GFP-labeled *Tdc2* cells. GFP-positive cells inside the red lines were collected for sequencing. **d**, Number of cells used in the sequencing analysis. **e**, Co-clustering frequency matrix from the iterative clustering analysis with 100 random samplings. The plot shows the probability of co-occurrence in the same cluster for given pairs of cells. **f-i**, tSNE plots of sequenced *Tdc2* cells, color-coded for the *nvy* locus genotypes (**f**; wild-type in black, Δ*nvy* in white), cell clusters (**g**), and expression levels of *Tdc2* (**h**) or *nvy* (**i**). **j**, Histogram of *Tdc2* cells according to the expression level of *nvy*. **k**, Ratio of *nvy*-expressing cells within each cluster. Red intensity corresponds to the level of *nvy* expression shown in **i-j**. Total cell numbers for each cluster are shown at the center. **l**, A volcano plot of DEGs analyzed in all *Tdc2* cells. Dots are plotted according to the fold change (FC) and the p-value (by Mann-Whitney U-test) of each gene when the Δ*nvy* mutant cells were compared against the wild-type cells.

**Extended Data Fig. 9.**
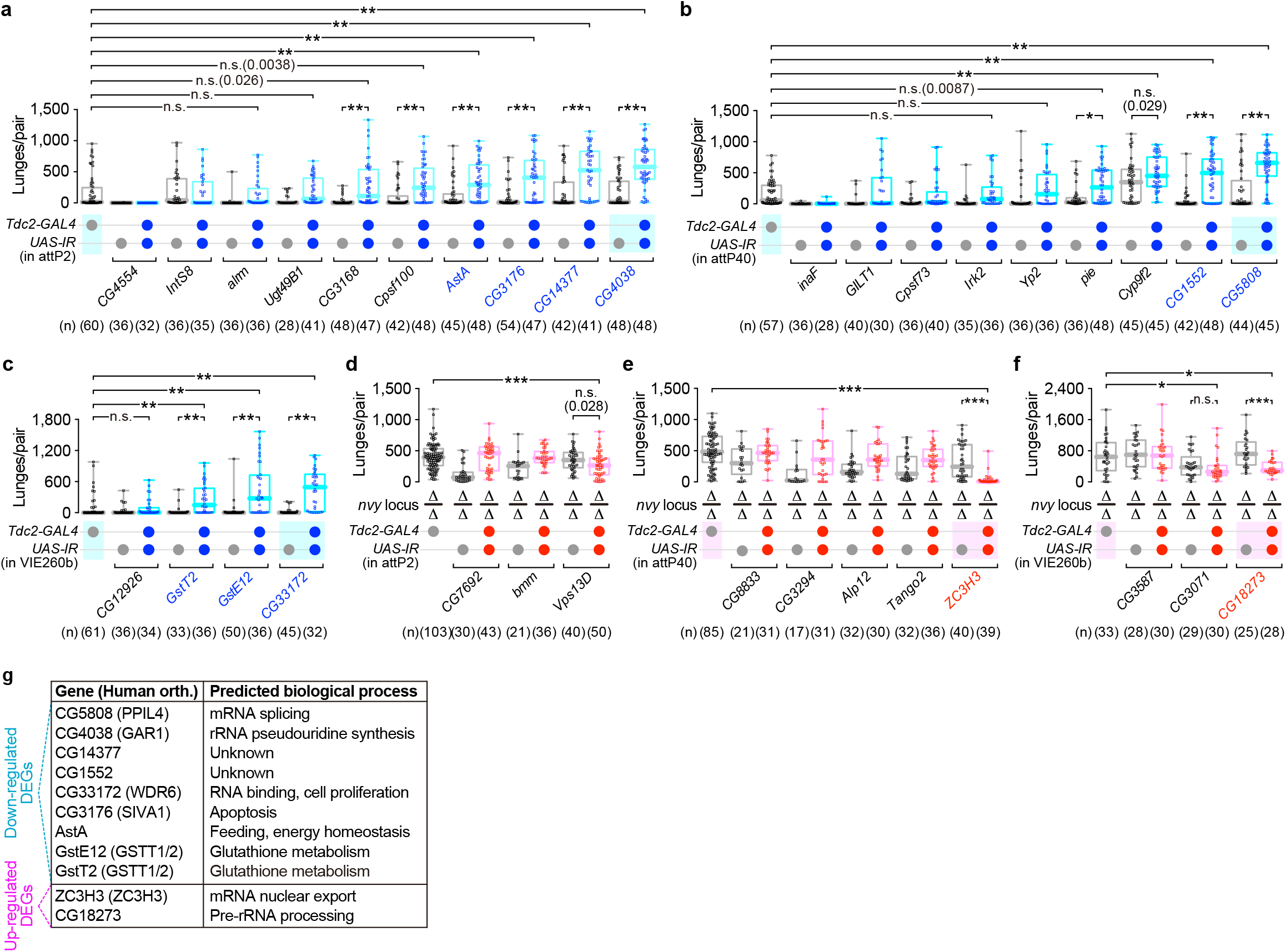
Aggressive behaviors by males following *Tdc2-GAL4*-driven RNAi of DEGs found in cluster #5. **a-c**, Lunges performed by males with *Tdc2-GAL4* driving RNAi constructs of down-regulated genes found in cluster #5. *UAS-IR* constructs were inserted either in attP2 (**a**), attP40 (**b**), or VIE260b (**c**). ** p < 0.01, * p < 0.05, n.s. p ≥ 0.05 (Kruskal-Wallis one-way ANOVA and post-hoc Mann-Whitney U-test with Bonferroni correction). **d-f**, Lunges performed by Δ*nvy* males with *Tdc2-GAL4* driving RNAi constructs of up-regulated genes found in cluster #5. All tested strains (*GAL4*-only, *UAS-IR*-only, and the knockdown mutants harboring both *GAL4* and *UAS-IR*) were made in the homozygous Δ*nvy* background. *UAS-IR* constructs were inserted either in attP2 (**d**), attP40 (**e**), or VIE260b (**f**). *** p < 0.0005, * p < 0.05, n.s. p ≥ 0.05 (Kruskal-Wallis one-way ANOVA and post-hoc Mann-Whitney U-test with Bonferroni correction). Knockdown of genes written in blue or red letters showed significant changes in lunges compared to both genetic controls. Note that data with shaded boxes are re-plotted in Fig. 5d and f. **g**, Predicted biological processes based on FlyBase for DEGs in cluster #5 that showed behavioral phenotypes in the RNAi experiments. Human orthologs are shown in parentheses.

## Materials and Methods

### Fly strains

#### Origins of fly lines

Full genotypes of flies used in experiments are listed in Supplementary Table S4. Canton-S originally from the lab of Dr. Martin Heisenberg (University of Wurzburg) was used as the wild-type strain. *UAS-IR* lines used in the pan-neuronal RNAi screen (Supplementary Tables S1 and S2) were selected from the KK collection in Vienna *Drosophila* Resource Center (VDRC), including *UAS-IR-nvy* (KK107374; VDRC #100273, RRID:FlyBase_FBst0472147) used in Fig. 1b and Extended Data Fig. 1b. Other *UAS-IR* lines were obtained from the TRiP collection in Bloomington *Drosophila* Stock Center (BDSC; University of Indiana), including another *UAS-IR-nvy* (JF03349; RRID:BDSC_29413) used in Extended Data Fig. 1a. The Exelixis deficiency *Df(2R)Exel6082* was obtained from BDSC (RRID:BDSC_7561). The following GAL4 lines were obtained from BDSC: *Akh* (RRID:BDSC_25684), *AstA*^1^ (RRID:BDSC_51978), *AstA*^2^ (RRID:BDSC_51977), *AstC* (RRID:BDSC_52017), *Burs* (RRID:BDSC_51980), *Capa* (RRID:BDSC_51969), *Crz* (RRID:BDSC_51975), *Dh31* (RRID:BDSC_51988), *Dh44* (RRID:BDSC_51987), *Dsk* (RRID:BDSC_51981), *ETH* (RRID:BDSC_51982), *FMRFa* (RRID:BDSC_51990), *Mip* (RRID:BDSC_51983), *NPF* (III) (RRID:BDSC_25682), *Pdf* (II) (RRID:BDSC_6900), *Proc* (RRID:BDSC_51971), *amon* (RRID:BDSC_30554), *ato*^10^ (RRID:BDSC_9494), *ato^14a^* (RRID:BDSC_6480), *ey^3–8^* (RRID:BDSC_5534), *ey^4–8^* (RRID:BDSC_5535), *GH146* (RRID:BDSC_30026), *Orco^11.17^* (RRID:BDSC_26818), *Poxn^1–7^* (RRID:BDSC_66685), *Ddc^4.3D^* (RRID:BDSC_7010), *Ddc^4.36^* (RRID:BDSC_7009), *Tdc2* (RRID:BDSC_9313), *Trh* (RRID:BDSC_38389), 5-HT1B (II) (RRID:BDSC_27636), 5-HT1B (III) (RRID:BDSC_27637), R11H09 (RRID:BDSC_48478), R15F02 (RRID:BDSC_48698), R16F12 (RRID:BDSC_48739), R17C11 (RRID:BDSC_48763), R20G01 (RRID:BDSC_48611), R27G01 (RRID:BDSC_49233), R38G08 (RRID:BDSC_50020), R70B01 (RRID:BDSC_39511), R84H09 (RRID:BDSC_47803), R93G12 (RRID:BDSC_40667). *8XLexAop2-FLPL* (in attP40) was obtained from BDSC (RRID:BDSC_55820). *UAS-Dicer2* (X) from BDSC (RRID:BDSC_24644) was used in the *w^+^* background. *fru^GAL4^* (described in Stockinger, *et al*.^52^), and *ppk23-GAL4* (described in Toda, *et al*.^53^) were gifts from Dr. Barry Dickson (HHMI Janelia Research Campus). *dsx^GAL4^*, originally described in Rideout, *et al*.^54^, was a gift from Dr. Stephen Goodwin (University of Oxford). *NP2631*, characterized in Yu, *et al*.^55^, was a gift from Dr. Daisuke Yamamoto (Tohoku University).*ppk25-GAL4*, originally described in Starostina, *et al*.^56^, was kindly shared by Dr. David Anderson (California Institute of Technology). *elav-GAL4* (III) was originally described in Luo, *et al*.^57^ and was used in Yapici, *et al*.^58^ 10XUAS-IVS-mCD8::GFP (in VK00005) and *20XUAS-IVS-Syn21-GFP-p10* (in attP2), originally described in Pfeiffer, *et al.^59^*, were created by Dr. Barret Pfeiffer in the lab of Dr. Gerald Rubin (HHMI Janelia Research Campus) and kindly shared by Dr. David Anderson. *20XUAS-IVS-Syn21-CsChrimson::tdTomato3.1* (in attP2), used in Watanabe, *et al*.^60^, was created by Dr. Barret Pfeiffer in the lab of Dr. Gerald Rubin and kindly shared by Dr. David Anderson. *10XUAS-IVS-Kir2.1^eGFP^* (in attP2), described in von Reyn, *et al*.^61^, was a gift from Dr. David Anderson. *Tub-FRT-GAL80-FRT*, originally described in Gordon, *et al*.^62^, was a gift from Dr. Kristin Scott (University of California, Berkeley). *Tub-FRT-stop-FRT-GAL80*, described in Bohm, *et al*.^63^, was a gift from Dr. Bing Zhang (University of Missouri). *hs-Cre* (X) was a gift from Dr. Konrad Basler (University of Zurich).

#### Genetic intersection labeling nvy-positive or -negative Tdc2 neurons

Genetic access to each subpopulation shown in Fig. 3 was achieved by expression of GAL80, an inhibitor of GAL4 initially utilized in *Drosophila* by Lee and Luo^64^, in undesired areas.

*nvy*(+) *Tdc2(+)*:

*w; Tdc2-GAL4, nvy^LexA^/Tub-FRT-GAL80-FRT, 8XLexAop2-FLPL; 10XUAS-IVS-XX/+*

In these flies, GAL80 is ubiquitously expressed under the tubulin promoter. Cells labeled by *nvy^LexA^* express flippase which excises the *GAL80* coding sequence flanked by flippase recognition targets (FRTs). This allows *Tdc2-GAL4-driven* expression of effectors (XX: GFP or Kir2.1^eGFP^) selectively in *nvy*-positive cells.

*nvy*(-) *Tdc2*(+):

*w*; *Tdc2-GAL4*, *nvy^LexA^*/*Tub-FRT-stop-FRT-GAL80*, *8XLexAop2-FLPL*; *10XUAS-IVS-XX*/+

As a transcriptional stop cassette flanked by FRTs precedes the *GAL80* coding sequence*, nvy*-positive *Tdc2* cells flip out the stop cassette, leading to GAL80-dependent suppression of GAL4. The remaining *Tdc2* cells, namely the *nvy*-negative *Tdc2* subpopulation, can express the effector.

### Generation of transgenic lines

*UAS-nvy*, NHR domain-deleted versions of *UAS-nvy, LexAop2-nvy, UAS-hMTG8*, and *UAS-hMTG16* lines were generated by ΦC31 integrase-mediated transgenesis as previously described^65^. Primer sequences are listed in Supplementary Table S5.

The *nvy* CDS (2,232 bp) was amplified from cDNA of the Canton-S strain. The CDS confirmed by sequencing is shown in Supplementary Information. Either three tandem c-Myc (3xMyc; EQKLISEEDLEQKLISEEDLEQKLISEEDL) or HA (3xHA; YPYDVPDYAGYPYDVPDYAGSYPYDVPDYA) epitope tag was attached to the 5’ end of the *nvy* CDS. As for the domain-deletion mutants of *nvy*, primers were designed to skip each NHR region (NHR1, 631–924th; NHR2, 1,360–1,440th; NHR3, 1,540–1,686th; NHR4, 1,777–1,890th nucleotides within the *nvy* CDS).

The original CDSs of the human *MTG8b* (1,815 bp; GenBank: D14821.1) and *MTG16b* (1,704 bp; GenBank: AB010420.1) genes were codon-optimized for expression in *Drosophila* (nucleotide sequences shown in Supplementary Information) by GenScript UAS Inc. (Piscataway, NJ).

To make the *10XUAS* constructs, the backbone plasmid pJFRC-MUH (RRID:Addgene_26213) was inserted with the intervening sequence (IVS)^66^ downstream of the hsp70 promoter, between *BglII* and *NotI* sites. As for the *13XLexAop2* constructs, pJFRC48-13XLexAop2-myr::tdTomato (a derivative of pJFRC19-13XLexAop2-myr::GFP (RRID:Addgene_26224) originally created by Pfeiffer, *et al*.^66^) was used as the backbone plasmid. Linker sequences containing the *Not*I site and the Kozak sequence (CAAA) were added right upstream to each CDS. Fragments and vectors were digested with *Not*I/*Xba*I, followed by ligation using T4 DNA ligase (NEB #M0202). Integrities of the resulting plasmids were confirmed by DNA sequencing. Plasmids were targeted to the attP site at VK00005 (RRID:BDSC_9725) using ΦC31 integrase-mediated transgenesis by BestGene Inc. (Chino Hills, CA). Transformants were selected by the eye color marker, and the presence of inserted CDSs were confirmed by PCR genotyping. All transgenic lines were backcrossed to the wild-type Canton-S for 6 generations prior to experiments.

### CRISPR/Cas9-mediated generation of *nvy* mutants

Δ*nvy* and *nvy_LexA_* lines were created based on the CRISPR/Cas9-mediated genome editing^67^ as follows. Primer sequences are provided in Supplementary Table S5.

Target sites for guide RNAs were searched by using the online CRISPR Target Finder available at the flyCRISPR website (http://flycrispr.molbio.wisc.edu/) with default settings. Within the genome region surrounding the 1st exon of the *nervy* gene (Dmel\CG3385), the following sites with no detectable off-targets were selected (PAM sequence underlined):

gRNA target #1: 5’-TGATGTTTTCGTCTATCGCC<u>CGG</u>-3’
gRNA target #2: 5’-TCATTGTTTGGAACTATAAT<u>AGG</u>-3’

Primers containing linkers attached to each target without the PAM sequence were used for PCR with pCFD4 (RRID:Addgene_49411) as a template. The amplified 598-bp fragment was ligated with *Bbs*I-digested pCFD4, and the resulting pCFD4-nvy-gRNA-1 plasmid was injected into embryos of the *vas-Cas9* (X) strain (RRID:BDSC_51323). The F0 adults (17 males and 17 females) were crossed individually with a balancer line, and F1 flies (5–12 males and 5 females from each F0 cross) were screened by PCR genotyping. Among 354 F1 individuals, two sibling lines with an identical 513-bp deletion (from −132 bp to +134 bp of the 1st exon) were found, designated herein as Δ*nvy*. The Δ*nvy* line was backcrossed to the wild-type Canton-S for 11 generations prior to experiments.

Our initial attempt for knock-in line generation using the above gRNA plasmid failed (none out of 478 F1 individuals from 16 F0 crosses were DsRed-positive) presumably due to low effectiveness in genome editing. To overcome this issue, we prepared a secondary plasmid (pCFD4-nvy-gRNA-2) for additional supply of gRNAs that target the following sites:

gRNA target #3: 5’-GTTTCCAAGTTCCCAGGTTC<u>CGG</u>-3’
gRNA target #4: 5’-CACCAACAACACAACATCGG<u>CGG</u>-3’

To construct the *nvy^LexA^* knock-in plasmid, pHD-DsRed (RRID:Addgene_51434) was used as backbone. Left (from −1,688 to −1 bp of the 1st exon) and right (from +140 to +1,859 bp of the 1st exon) homologous arms were amplified from genome DNA of the *vas-Cas9* (X) strain. Point mutations were introduced within the PAM sequences of gRNA target sites to avoid plasmid cleaving by Cas9. The *nls::LexA::p65* CDS was amplified from pBPnlsLexA::p65Uw (RRID:Addgene_26230). The 1st exon of *nvy*, of which the start codon was mutated from ATG to TAG, followed by the 139-bp downstream region was amplified from the genome DNA. The knock-in plasmid was constructed by using In-Fusion HD Cloning kit (Takara Bio USA #639650) or NEBuilder HiFi DNA Assembly kit (New England Biolabs #M5520) in two steps: the left homologous arm, *nls::LexA::p65* CDS, and the 1st exon of *nvy* were first fused with the *Xho*I/*Spe*I-digested pHD-DsRed vector; then the resulting plasmid was digested with *Not*I/*Eco*RI followed by insertion of the homologous right arm to generate pHD-DsRed-nvy-LexA.

Three plasmids (pCFD4-nvy-gRNA-1, pCFD4-nvy-gRNA-2, and pHD-DsRed-nvy-LexA) were co-injected to embryos of the *vas-Cas9* (X) strain. The F0 adults (24 males) were crossed each with the balancer line, and F1 offspring were screened for DsRed expression in compound eyes under a fluorescent microscope. From 382 F1 males collected from 7 F0 crosses, 15 individuals were found positive for DsRed. Insertion of *LexA* was confirmed by genotyping PCR with the primers used for the pHD-DsRed-nvy-LexA plasmid construction. Three candidate lines were backcrossed to the wild-type Canton-S for 6 generations, and the *DsRed* sequence flanked by two *loxP* sites was excised by crossing with *hs-Cre* (X). Genomic regions surrounding the 1st exon of *nvy* were analyzed by Southern blot as described below. One of the validated alleles was used as *nvy^LexA^* for further experiments.

The nucleotide sequences of generated plasmids are provided as Supplementary files. Plasmid injections to fly embryos were performed by BestGene Inc.

### Southern blot

Two hundred adult flies per genotype were grinded in 800 μL of TE buffer (Tris/HCl (pH 9), 100 mM EDTA) supplemented with 1% SDS, followed by incubation at 65°C for 30 min. Samples were added with 300 μL of 3 M potassium acetate and placed on ice for 30 min. After centrifugation at 13,000 rpm for 20 min at 4°C, the supernatant (600 μL) was collected and mixed with a half volume of isopropanol. Samples were centrifuged at 13,000 rpm for 10 min, and the pellet was washed with 70% ethanol. Precipitates were dried and dissolved in 500 μL of TE buffer. Samples were then treated with RNase A (0.4–0.8 mg/mL) at 37°C for 15 min. For purification, each sample was mixed vigorously with the same volume of PCI (phenol:chloroform:isoamyl alcohol = 25:24:1, v/v). After centrifugation at 13,000 rpm at 5 min, the aqueous upper layer was collected and mixed vigorously with the same volume of chloroform, followed by another centrifugation. The upper layer (400 μL) was further subjected to ethanol precipitation. The final precipitates obtained were dried and dissolved in 100 μL of TE buffer. The typical yield of genomic DNA extracted from 200 flies was 0.2–0.5 mg.

Ten to twenty micrograms of genomic DNA per each genotype was digested with *HindIII* at 37°C for overnight. Electrophoresis was performed using a 0.7% agarose gel. Digoxigenin (DIG)-labeled DNA molecular weight maker III (Roche #11218603910) was loaded as a marker. The gel placed on a shaker was sequentially subjected to depurination (in 0.25 N HCl for 10 min), denaturation (in 0.5 M NaOH, 1.5 M NaCl for 15 min x 2), neutralization (in 0.5 M Tris/HCl (pH 7.5), 1.5 M NaCl for 15 min x 2), and equilibration (in 20 x SSC for 10 min). DNA was transferred to a nylon membrane (Roche #1120929901) for overnight, by sandwiching between paper towels soaked in 20 x SSC under a weight of 1.5 kg. DNA was immobilized onto the membrane by using UV Stratalinker 2400 (Stratagene).

DIG-labeled DNA probes were synthesized using PCR DIG Probe Synthesis Kit (Roche #11636090910). Primers were designed to target either external (676 bp; the genomic region from −1,986 to −2,661 bp upstream of the *nvy* exon 1) or internal (621 bp; 660–1,280th nucleotides within the *nls::LexA::p65* CDS) regions of the *LexA* knock-in construct, as shown in Supplementary Table S5. The DIG-labeled probes were hybridized to the membrane in DIG Easy Hyb hybridization buffer (Roche #11603558001) at 49°C for overnight. The membrane was sequentially washed with a low stringency buffer (2 x SSC, 0.1% SDS) at room temperature for 5 min x 2, and with a pre-warmed high stringency buffer (5 x SSC, 0.1% SDS) at 68°C for 15 min x 2. After another brief wash with a buffer (from DIG Easy Hyb kit), the membrane was soaked in a blocking buffer (from DIG Easy Hyb kit) at 4°C for overnight. Alkaline phosphatase-conjugated anti-DIG Fab fragments (Roche #11093274910, RRID:AB_514497) were freshly added to the blocking buffer at 1:10,000, and the membrane was incubated at room temperature for 30 min. The membrane was washed with the wash buffer for 15 min x 2, followed by a brief equilibration in a detection buffer (from DIG Easy Hyb kit). As a chemiluminescence substrate, CDP-Star (Roche #11759051001) was freshly diluted to 1:200 in the same buffer. Signals were developed on autoradiography films (Genesee Scientific #30-507).

### Western blot

Sixty to ninety adult flies (5–7 days post eclosion) were snap-frozen in liquid nitrogen. The fly heads were separated from other body parts in liquid-nitrogen chilled metal sieves. Collected heads were grinded in 60–90 μL of ice-cold extraction buffer (20 mM HEPES (pH 7.5), 100 mM KCl, 10 mM EDTA, 0.1% Triton X-100, 1 mM DTT, 5% glycerol; according to Thomas, *et al*.^68^) with disposable pestles, followed by centrifugation at 1,600 x g for 20 min at 4°C. The supernatant was mixed with 4 x Laemmli Sample Buffer (Bio-Rad #1610747), and samples were heated in boiling water for 5 min.

Proteins were separated in 4–20% Mini-PROTEAN TGX Precast Protein Gels (Bio-Rad #4561096) and transferred to 0.45 μm pore-size nitrocellulose membranes (Bio-Rad #1620215). Membranes were shaken in TBST (20 mM Tris/HCl (pH 7.6), 150 mM NaCl, 0.1% Tween-20) supplemented with 5% blotting-grade blocker (Bio-Rad #1706404) at room temperature for 2–3 h. After washing in TBST for 10 min x 3, membranes were incubated with primary antibodies (1:1,000–10,000 dilution in 2–5% skim milk/TBST or Can Get Signal solution 1 (Toyobo #NKB-201)) at room temperature for 1–2 h. Membranes were washed in TBST for 10 min x 3, followed by reaction with horseradish peroxidase (HRP)-conjugated secondary antibodies (1:10,000 dilution in 2–5% skim milk/TBST or Can Get Signal solution 2 (Toyobo #NKB-301)) at room temperature for 1-2 h. After the final wash in TBST for 10 min x 3, membranes were treated with Clarity Western ECL Substrate (Bio-Rad #1705061). Signals were developed on autoradiography films (Genesee Scientific #30-507).

Detailed information for antibodies and incubation conditions are provided in Supplementary Table S6.

### Immunoprecipitation

Immunoprecipitation of Myc- and HA-tagged Nvy proteins were preformed essentially as described previously^69^. Tagged Nvy proteins were pan-neuronally expressed under the control of *elav-GAL4*. Heads from 100–120 flies were isolated using liquid-nitrogen chilled metal sieves as described above, followed by homogenization in 700 μL of buffer B (20 mM Tris/HCl (pH 7.6), 150 mM NaCl, 5 mM MgCl_2_, 10% sucrose, 1% glycerol, 1 mM EDTA, protease inhibitors (1 tablet of cOmplete™ Protease Inhibitor Cocktail (Roche #11697498001) dissolved in 50 mL)) supplemented with 1% CHAPS. Homogenates were first centrifuged at 16,000 x g for 30 min at 4°C, and the supernatants were centrifuged again at 16,000 x g for 20 min at 4°C. Cleared lysates (650 μL) were collected carefully using capillary pipet tips. Lysates were separated into three groups of 200 μL each and added with 800 μL of buffer A (20 mM Tris/HCl (pH 7.6), 150 mM NaCl, 1 mM dithiothreitol, 3 mM MgCl_2_, 1 mM EGTA). The remaining lysates were stored at −20°C to be used as “inputs”.

Protein G PLUS-Agarose (Santa Cruz Biotechnology #sc-2002, RRID:AB_10200697) was washed with buffer A, and 10 μL of 50% bead slurry was added to each sample. As a pre-cleaning step, samples were gently rotated for 1 h at 4°C. Samples were then centrifuged at 1,000 x g for 30 s at 4°C, and collected supernatants were centrifuged again at 3,000 x g for 30 s at 4°C. For each genotype, one sample was kept as negative control without antibody, another sample was added with 2.5 μL of normal rat IgG (0.4 mg/mL; Santa Cruz Biotechnology #sc-2026, RRID:AB_737202), and the last sample was added with 1 μL of anti-c-Myc rat IgG1 (1 mg/mL; clone JAC6, Abcam #ab10910, RRID: AB_297569). The antibody binding was performed for 2–3 h at 4°C on a rotator. To prepare the beads for precipitation, Protein G PLUS-Agarose was washed and suspended in buffer B supplemented with 0.2% CHAPS and 1% bovine serum albumin, and incubated for 30 min at 4°C on a rotator. Beads were washed twice in buffer B, and then suspended to make 50% slurry. For immunoprecipitation, each sample was added with 40 μL of bead slurry and incubated for overnight at 4°C on a rotator. After centrifugation at 1,000 x g for 30 s at 4°C, precipitated samples were washed twice with 0.5 mL of buffer A supplemented with 0.2% CHAPS. The final precipitates were suspended in 20 μL of 2 x Laemmli buffer and heated in boiling water for 10 min. Western blot was performed as described above.

### Immunohistochemistry

Immunohistochemistry of fly brains essentially followed the method described by Van Vactor, *et al*.^70^. Fly brains were dissected in PBS, and then incubated in the fixing solution (2% formaldehyde, 75 mM L-lysine in PBS) at room temperature for 1–1.5 h. All reactions from fixation to clearing were carried out in a well of 6 x 10 microwell minitray (Thermo Fisher Scientific #439225). Brains were washed in PBST (0.3% TritonX-100 in PBS) for 5 min x 3, followed by incubation in a blocking solution (5% heat-inactivated normal goat serum, 0.3% TritonX-100 in PBS) for 30 min. Primary antibodies diluted with the blocking solution (1:10 or 100 for mouse anti-BRP (Developmental Studies Hybridoma Bank nc82 (supernatant or concentrated), RRID: AB_2314866), 1:1,000 for chicken anti-GFP (Abcam #ab13970, RRID: AB_300798), 1:1,000 for rabbit anti-DsRed (Takara Bio USA #632496, RRID: AB_10013483)) were applied to the samples at 4°C for 2 days. The brains were washed in PBST for 10 min x 3, and then incubated in secondary antibodies diluted with the blocking solution (1:100 for goat anti-mouse Alexa 633 (Thermo Fisher Scientific #A-21052, RRID: AB_2535719), 1:100 for goat anti-chicken Alexa 488 (Thermo Fisher Scientific #A-11039, RRID: AB_2534096), 1:100 for goat anti-rabbit Alexa 568 (Thermo Fisher Scientific #A-11036, RRID: AB_10563566)) at 4°C for overnight. Brains were washed in PBST for 10 min x 3, and then incubated in the clearing solution (50% glycerol/PBS) at room temperature for 2 h. Samples were mounted in Vectashield (Vector Laboratories, #H-1000) onto a slide glass. Images were acquired by FV-3000 confocal microscopy (Olympus America; kindly shared by Dr. Samuel Pfaff at Salk Institute). Stacked images of maximum z-projections were generated on Fiji software^71^ (RRID: SCR_002285; https://fiji.sc/).

### Social behavior experiments

#### Behavioral apparatus

Twelve-well acrylic chambers were designed as previously described^9^. Each arena had a diameter of 16 mm and a height of 10 mm. The entire floor was covered with apple juice gel (Minute Maid 100% apple juice, 2.25% agarose, 2.5% sucrose w/v) as food source. The inner wall and ceiling were coated with Insect-A-Slip (BioQuip Products #2871C) and Surfasil Siliconizing Fluid (Thermo Fisher Scientific #TS-42800), respectively.

To allow flies to perform wider repertoire of behaviors during inter-male encounters, chambers with larger space and limited food source, similar to those described previously^20,60^, were used in Fig. 2a-d and Extended Data Fig. 3. Each arena had a size of 40 mm x 50 mm and a height of 70 mm. A small window for fly introduction was made at 40 mm height from the bottom. The apple juice gel was poured into a depressed area of 10 mm x 10 mm with a depth of 5 mm located at the center of each arena. A transparent acrylic plate was placed on top of the chamber. The inner wall and ceiling were coated as above.

The chambers were lit from underneath by LED backlights. For optogenetic experiments, 850-nm infrared backlights (Sobel Imaging Systems #SOBL-150×100-850) were used instead. Movies were taken using the Point Grey Flea3 USB3.0 digital cameras (FLIR #FL3-U3-13Y3M-C) controlled by the BIAS acquisition software (IORodeo; https://bitbucket.org/iorodeo/bias). The camera was mounted with a machine vision lens (Fujinon #HF35HA-1B). For optogenetic experiments with the infrared backlights, an infrared longpass filter (Midwest Optical Systems #LP780-25.5) was attached to the camera. Recording was performed either at 30 fps for the 1st round of RNAi screen (Fig. 1a; left), or at 60 fps for the rest of all experiments. The optogenetic stimulation was performed using 655-nm red light LEDs controlled by the Arduino Uno board (Arduino) with a custom program as described previously^72^.

#### Fly preparation and behavioral assays

Parental flies (no more than 20 females and 10 males per bottle) were reared on 50 mL of standard cornmeal-based food, and were transferred to fresh food every 2–3 days. The small-body sized flies were obtained as previously described^9^, by rearing offspring from a larger group of parental flies (70 females and 30 males) that laid eggs on less amount of food (5 mL) for 1–2 days. Offspring flies were collected on the day of eclosion into vials with standard fly food medium. Adult males and females were kept separately to avoid mating, except when mated females were prepared for targets in some experiments. For optogenetic experiments, adult testers were reared on food supplied with 0.2 mM all-*trans* retinal (Sigma-Aldrich #R2500, 20 mM stock solution prepared in 95% ethanol), and the vials were covered with aluminum foil to avoid light exposure. Flies were kept either as a group of up to 15 (“group-reared”) or one (“single-reared”) per vial at 25°C with 60% relative humidity, in a 12-h light/dark cycle (light phase 9AM–9PM). Flies were transferred to new vials with fresh food after every 3 days. To keep track of each fly’s identity within a pair of males with different genotypes, the tip of either one of the wings were clipped by a razor under brief CO_2_ anesthesia. This marking treatment itself does not reduce the level of lunge or wing extension by males under our experimental conditions^39^.

Behavior experiments were performed in the evening (4–9PM) at 22–25°C. When pairing group-reared flies of same genotypes, two flies were always taken from different vials to make a pair that has never met each other during their adulthood. For male-male and female-female pairs tested in the 12-well chamber, flies were introduced by gentle aspiration and acclimated for 5 min prior to the 30-min recording. In case of male-female pairs, females were first introduced into the arenas, and males were trapped between two small plastic tips set upon the lid to prevent contact with females. After 5 min of acclimation, all males were simultaneously introduced to the arenas by sliding the lid, and the 1-h recording was immediately started. In competitive copulation assays, male pairs were first loaded into the 12-well chamber, and one virgin female was trapped upon each well as described above. After acclimating the male pairs for 15 min, the females were simultaneously introduced to all arenas, and recording was immediately started. For experiments using the large chamber where we aimed to capture the flies’ behaviors from the first encounter, the recording was started before introduction of flies to the arenas, and the movie was taken for 35 min. The 30-min time window after the entrance of the second fly to the arena was used for behavior analysis.

Courtship memory assay was performed essentially as described previously^73^. In brief, a sexually naïve male was introduced into a 1.5 mL tube with food, and either kept alone as the sham-trained group or paired with an unreceptive mated female as the trained group. Females were removed after 5 h of training, and the males were left in the same tubes for 1 h. Males were then transferred into the 12-well chamber, and a new set of wild-type mated females was loaded as above. The 30-min recording was started immediately after the females were introduced into the arenas.

#### Behavioral quantification

The 30-fps movies recorded in the 1st round of RNAi screen (Fig. 1a, left; Supplementary Table S1) were analyzed by CADABRA software^74^ on MATLAB (The Mathworks). The program was slightly modified to make it compatible with later MATLAB versions (2014b and 2019a) without affecting its functionality. Flies were tracked using the “qtrak” function. Lunges were detected using the analysis program that accompanies CADABRA, by applying the parameters originally described by Dankert, *et al*.^74^ The radius of the circular region of interest was set to 6.5 mm, which is approx. one fly body length smaller than the actual well size (8 mm radius), to exclude movements of flies staying close to or climbing on the wall that may lead to false positive detections.

The 60-fps movies recorded for the rest of all experiments were processed by the FlyTracker program^75^ (version 1.0.5) on MATLAB2014b. For pairs of flies with different conditions (sexes, genotypes, and/or rearing conditions), the identities of tester and target flies (marked by the clipped wing) were manually corrected throughout the movie. Behaviors were quantified using automated classifiers based on the machine-learning system JAABA^76^. The classifiers for lunges, headbutts, and wing extensions have been used in our recent studies^39,40^, and that for wing threat was newly created here. The details in development and performance of classifiers will be described elsewhere (manuscript in preparation). As a post-processing step to remove false-positives, extremely short bouts (< 50 ms for lunges and headbutts, and < 100 ms for wing extensions) were omitted from quantification. The time of one fly orienting the other (“time orienting”) was previously defined^39^ as the duration in which the following conditions are met: (1) the target fly is heading towards the target fly (within ± 60° of facing angle), (2) two flies are in close proximity (within 5 mm of distance), and (3) the target fly is moving (above 0.1 mm/s of velocity). Note that this orienting includes “chasing” which has been observed in the context of both courtship and aggression^74^.

In general, our lunge classifier has been trained to detect lunge bouts with a reasonably high precision (89%) and recall (88%). When testing male-female pairs, however, even a few cases of false-positives due to tracking errors might significantly affect the statistics of rarely observed male-to-female lunges. For this reason, we manually confirmed all male-to-female lunges detected in Extended Data Fig. 4b-c. As a result, 45 or 956 total bouts originally detected in either the wild-type (n = 36) or Δ*nvy* (n = 35) males were manually confirmed to contain 1 or 928 true lunges, respectively (Extended Data Fig. 4b).

Statistical analyses for RNAi screen were performed on MATLAB2014b (“ranksum”). RNAi mutants that (1) passed the Benjamini-Hochberg FDR test^77^ of 0.05 and (2) showed the median lunge numbers (per pair in 30 min) more than 3 were selected as hits. Statistical analyses for the rest of all experiments were carried out using Prism 6 (GraphPad Software). Multiple comparisons among different genotypes were performed using the Kruskal-Wallis test followed by the post-hoc Mann-Whitney U-test. When comparing paired datasets among different genotypes within a pair of flies or optogenetic stimulation periods within the same fly groups, the post-hoc Mann-Whitney signed rank test was used. Bonferroni correction was applied to adjust the p-values.

#### Behavioral feature analysis

The two-dimensional heat-map represents a kernel density estimate of the joint distribution of two kinematic features. Frame-wise values of facing angles (“facing_angle”) and distance between two flies (“dist_to_other”) were extracted from-feat.mat files created by the FlyTracker program, and were pooled across flies with the same genotype/rearing conditions. For Fig. 2b, data points each associated with a lunge followed by the same fly’s next lunge with an interval shorter than 0.5 s were excluded.

The kernel density estimation at 90 points equally spaced along each feature dimension was performed through the “ksdensity” function on MATLAB, using a normal kernel with default parameters. The heat-map used the color code originally created by Dr. Jonathan Herman (University of California, Davis) available on Github (https://jdherman.github.io/colormap/) to represent the probability density at the estimated points within each panel. Facing angles were plotted in the range from 0 to π rad according to the design of tracking algorithm. For the distance between flies, the range from 1.0 to 15.0 mm was arbitrarily chosen, which covered 95.3% and 99.7% of total data points for wild-type and Δ*nvy* testers, respectively. The plot space was binned into 3 x 3 sections, and the percentage of event occurrence within each section was calculated. For one-dimensional features plotted on each axis, both Wilcoxon rank-sum test (for differences in data distribution based on median values) and Kolmogorov-Smirnov test (for differences in cumulative distribution functions) were used to compare the wild-type and Δ*nvy* testers. In addition, a two-dimensional Kolmogorov-Smirnov test^78^, using an implementation by Dr. Brian Lau (L’Institut du Cerveau et de la Moelle Épinière) available on Github (https://github.com/brian-lau/multdist), was applied to evaluate the difference between the two-dimensional distributions of features observed in “wild-type single-reared testers vs wild-type group-reared targets” and “Δ*nvy* testers vs wild-type group-reared targets”. P-values are summarized in Supplementary Information.

The scatter plot of lunge interval and maximum inter-lunge facing angle was generated using the feature data extracted above. Note that, unlike the two-dimensional heat-maps, all lunges with intervals shorter than 0.5 s were included for this analysis. The minimum measurable lunge interval was set to 1/60 s, as the movies were recorded at 60 fps. The plot space was arbitrary divided into three time-windows (< 2 s, 2–20 s, ≥ 20 s), and histograms of maximum inter-lunge facing angles were generated for each time section. The heights of histograms were normalized globally against the total number of lunges by each tester genotype. Maximum inter-lunge facing angles were plotted in 12 bins, each with a width of π/12. Statistical differences of histograms between the wild-type and Δ*nvy* testers were analyzed by the Kolmogorov-Smirnov test using the MATLAB built-in function “kstest2”.

### Single-cell RNA sequencing

#### Preparation of single-cell suspensions

Virgin male and female flies expressing mCD8::GFP under the control of *Tdc2-GAL4* in either wild-type *nvy* locus or homozygous Δ*nvy* mutant background were used. Adults were collected upon eclosion and kept as a group of 15 per vial for 5–7 days at 25°C, as done for social behavior experiments (see above).

Single-cell suspensions were prepared according to the protocol detailed in Li, *et al*.^79^ Fly brains were dissected and stored in ice-cold Schneider’s insect medium (Sigma-Aldrich #21720024) for up to 2 h. The brains were rinsed in cold RNase-free PBS and transferred to freshly made dissociation buffer (300 μL of 100 U/mL heat-activated papain (Worthington Biochemical Corporation #LK003178) added with 6 μL of 2.5 mg/mL liberase™ (Sigma-Aldrich #5401119001)), followed by an incubation for 20 min at 25°C under continuous shaking at 1,000 rpm. During this 20-min incubation, the suspension was pipetted for 30 times at the 5 and 10-min time points, and then forced through a 25G 5/8 needle for 7 times at the 15-min mark. One milliliter of ice-cold Schneider’s insect medium was added to terminate the enzymatic digestion. The suspension was then filtered through a cell strainer with 35 μm mesh size (BD Biosciences #352235) and centrifuged at 600 x g for 7 min at 4°C. The pellet was re-suspended in cold Schneider’s insect medium supplemented with 1 μg/mL DAPI (4’,6-diamidino-2-phenylindole; Thermo Fischer Scientific #D-1306). Samples were sorted using the BD Vantage DiVa™ Cell Sorter (BD Biosciences). Gates were set to collect viable (DAPI-negative) GFP-positive cells as shown in Extended Data Fig. 8c. Single cells were collected into individual wells of 96 well-PCR plates containing 9.5 μL/well of freshly made lysis buffer (provided in SMART-Seq v4 Ultra Low Input kit for Sequencing (Takara Bio USA #634893)). After the sorting, samples were immediately placed on dried ice and stored at −80°C until use. To prevent non-physiological transcriptional activities triggered during the single-cell preparation process, all solutions were supplemented with actinomycin D (Sigma-Aldrich #A1410) at the final concentration of 5 μg/mL.

In total, brains were dissected from 359 *nvy* wild-type (213 males and 146 females) and 381 Δ*nvy* (242 males and 139 females) flies in 10 experimental days. Lysates of 197 *nvy* wild-type (93 male- and 104 female-derived) and 216 Δ*nvy* (112 male- and 104 female-derived) cells were processed as below.

#### Single-cell sequencing

mRNA in the cell lysate was reverse-transcribed and amplified for 25 cycles using the SMART-Seq v4 Ultra Low Input kit for Sequencing (Takara Bio USA #634893) according to manufacturer’s instructions. To confirm the presence of GFP transcripts, each cDNA was subjected to PCR genotyping using Emerald AMP HS PCR Master Mix (Takara Bio USA #RR330B) and primers shown in Supplementary Table S5. As a result, 104 *nvy* wild-type (55 male- and 49 female-derived) and 114 Δ*nvy* (58 male- and 56 female-derived) GFP-positive samples were selected for sequencing. The amplified cDNAs were quantified by the Qubit^®^ 3.0 Fluorometer (Thermo Fischer Scientific #Q33216) and normalized to a concentration of 0.22 ng/μL. Sequencing libraries were prepared using the Nextera XT kit (Illumina #FC-131-1096) and mixed into 24 pools (12 samples per pool). After purification using the Agencourt AMPure XT beads (Beckman Coulter #A63881), the sample quality was checked with both the Qubit 3 Fluorometer and the High Sensitivity D1000 ScreenTape assay (Agilent Technologies #5067-5584). The libraries were equimolarly pooled, and the final concentration was estimated by qPCR using primers shown in Supplementary Table S5 and KAPA Library Quantification Kit Illumina Platforms KK4828 (Roche #07960166001) according to manufacturer’s instructions. Sequencing of 75 bp paired-end reads was performed with the Illumina NextSeq 500 sequencer.

#### Bioinformatics analysis

In total, 218 cells were sequenced. Reads were quality-tested using FASTQC^80^ and aligned to the *D. melanogaster* genome dm6 (from The FlyBase Consortium/Berkeley Drosophila Genome Project/Celera Genomics) using the alignment algorithm STAR^81^ (version 2.5.3a). Mapping was carried out using default parameters (up to 10 mismatches per read, and up to 9 multi-mapping locations) with additional code to filter out alignments that contain non-canonical junctions (--outFilterIntronMotifs RemoveNoncanonical). Raw gene expression was quantified using the software HOMER^82^ across exons, and the top isoform value was used to represent gene expression. Raw counts were processed using the supplied R^83^ script. In brief, cells containing the bottom 10% of raw sequence counts were filtered (22 cells), TMM normalization and size-factor correction was applied using the edgeR^84^ package (version 3.24.3). Then, the bottom 10% of cells with genes having normalized counts > 32 per cell were removed as cells with low gene expression (19 cells). As a summary, a sequence depth of 1 x 10^6^ reads per cell with 6 x 10^3^ average genes per cell (4,157 genes after the bottom 10% cut-off) was achieved for 171 cells (Extended Data Fig. 8d).

Expression values were log2-transformed, and tSNE^85^ was performed to generate the plots. The scrattch.hicat R package^50^ (version 1.0.0) was used to perform hierarchical iterative clustering^86^ on the normalized expression dataset (Supplementary Table S7). Default parameters were adjusted (see the supplied script), and a stochastic sampling and consensus clustering approach (run_consensus_clust) was used to assign cell cluster identity, as recommended by authors of the original code. Cell cluster co-occurrence was plotted with heatmap.2 from gplots in R. For differential expression analysis between the wild-type and Δ*nvy* mutant, genes with expression values of 0 in 75% or more of the cells were filtered out. Genes that met the following criteria were considered as DEGs: (1) p-values by Mann-Whitney U-tests lower than 0.05, (2) passed Benjamini-Hochberg FDR test of 0.2, and (3) fold change greater than 10 (|log_2_FC| > 3.321928095). For DEGs that showed behavioral phenotypes in the RNAi experiments, predicted biological processes and human orthologs were taken from the “Gene Ontology” and “Human Orthologs (via DIOPT v7.1)” sections in FlyBase (http://flybase.org/), respectively.

## Data availability

All data necessary to reproduce figure panels and statistical analyses, as well as other relevant files, are available upon request. The raw count table and the FASTQ flies are deposited to the Gene Expression Omnibus (GEO) database under the accession code GSE148630.

## Code availability

Codes necessary to reproduce the behavioral analyses are available on Github (https://github.com/asahinak/Ishii_etal_2020_behavioral_feature_analysis). R scripts necessary to reproduce the statistical analyses on sequencing data are provided as a supplementary file.

## Acknowledgements

We thank Drs. David Anderson, Gerald Rubin, and Barret Pfeiffer for sharing unpublished transgenic *Drosophila* strains with us; Drs. Stephen Goodwin, Daisuke Yamamoto, Kristin Scott, and Bing Zhang for other *Drosophila* strains; Dr. Samuel Pfaff for sharing the Olympus FV-1000 confocal microscopy with us, Drs. Christopher Kintner, Greg Lemke, Eiman Azim, and members of the Asahina lab for critical comments on the manuscript. The rabbit anti-Nvy antibody was kindly provided by Dr. Richard Mann (Columbia University). The antisera nc82 (anti-BRP), developed by Dr. Erich Buchner, were obtained from the Developmental Studies Hybridoma Bank, created by the NICHD of the NIH and maintained at The University of Iowa, Department of Biology, Iowa City, IA 52242. Stocks obtained from the Bloomington Drosophila Stock Center (NIH P40OD018537) were used in this study. Technical assistance for confocal imaging was provided by Drs. Uri Manor and Tong Zhang at the Waitt Advanced Biophotonics Core Facility of the Salk Institute, funded from NIH-NCI CCSG: P30 014195 and the Waitt Foundation. This work was supported by the Naito Foundation Grant for Studying Overseas to KI, JSPS Postdoctoral Fellowship for Research Abroad (28-869) to KI, and 1R35GM119844 from NIH/NIGMS to KA. The RNAi screen was partly conducted with support from Dr. David J. Anderson’s research group at California Institute of Technology.

